# Neuronal glucose metabolism sets cholinergic tone and controls thermo-regulated signaling at the neuromuscular junction

**DOI:** 10.1101/2021.09.28.462124

**Authors:** Yan Tang, Haihong Zong, Hyokjoon Kwon, Yunping Qiu, Jacob B. Pessin, Licheng Wu, Katherine A. Buddo, Ilya Boykov, Cameron A. Schmidt, Chien-Te Lin, P. Darrell Neufer, Gary J. Schwartz, Irwin J. Kurland, Jeffrey E. Pessin

**Affiliations:** Department of Medicine, Albert Einstein College of Medicine, Bronx, NY 10461, USA; Department of Neuroscience, Albert Einstein College of Medicine, Bronx, NY 10461, USA; Department of Molecular Pharmacology, Albert Einstein College of Medicine, Bronx, NY 10461, USA; The Fleischer Institute of Diabetes and Metabolism, Albert Einstein College of Medicine, Bronx, NY 10461, USA; East Carolina Diabetes and Obesity Institute and the Department of Physiology, Brody School of Medicine East Carolina University, Greenville, NC 27834, USA

**Keywords:** TIGAR, cholinergic neurons, acetylcholine, neuromuscular junction, skeletal muscle thermogenesis

## Abstract

Cholinergic and sympathetic counter-regulatory networks control numerous physiologic functions including learning/memory/cognition, stress responsiveness, blood pressure, heart rate and energy balance. As neurons primarily utilize glucose as their primary metabolic energy source, we generated mice with increased glycolysis in cholinergic neurons by specific deletion of the fructose-2,6-phosphatase protein TIGAR. Steady-state and stable isotope flux analyses demonstrated increased rates of glycolysis, acetyl-CoA production, acetylcholine levels and density of neuromuscular synaptic junction clusters with enhanced acetylcholine release. The increase in cholinergic signaling reduced blood pressure and heart rate with a remarkable resistance to cold-induced hypothermia. These data directly demonstrate that increased cholinergic signaling through the modulation of glycolysis has several metabolic benefits particularly to increase energy expenditure and heat production upon cold exposure.

**Highlights:**

1. Deficiency of a negative regulator of glycolysis (TIGAR) in cholinergic neurons increases the biosynthesis and content of the neurotransmitter acetylcholine.
2. Increased cholinergic tone reduces blood pressure and heart rate while enhancing signaling at neuromuscular junction.
3. Upregulation of neuromuscular junction activation provides protection against the paralytic curare and cold-induced hypothermia.
4. Modulation of cholinergic neuron glycolysis may provide a novel therapeutic approach for treatment of diseases stemming from reduced acetylcholine signaling such as myasthenia gravis and sarcopenic pre-synaptic dysfunction.

## Introduction

TIGAR (Tp53-induced Glycolysis and Apoptosis Regulator) was originally identified as a p53-inducible protein that functions as a fructose-2,6-bisphosphatase (F2,6P) but subsequently shown to have phosphatase activities for a variety of phosphorylated metabolic intermediates and allosteric regulators including 2,3-bisphospholgycerate (2,3BPG), 2-phosphoglycerate, phosphoglycolate and phosphoenolpyruvate (Bensaad, Tsuruta et al. 2006, Rigden 2008, Bolanos 2014, Tang, Chen et al. 2021). Due to its F2,6P dephosphorylation activity, TIGAR is generally considered as a suppressor of glycolysis as F2,6P is a potent allosteric activator for 6-phosphofructo-1-kinase (PFK1), the rate-limiting step in glycolysis. However, the dephosphorylation of 2,3BPG to generate 3-phosphoglycerate would also be expected to increase glycolysis (Bolanos 2014). In addition, the role of TIGAR in regulating carbohydrate metabolism is also complicated by the presence of the related phosphofructokinase bis-phosphatase (PFKBP) family that has both F6P 2-kinase as well as F2,6P bisphosphatase activities (Mor, Cheung et al. 2011). Thus, the biologic readout of TIGAR function is likely to be highly cell context dependent. In this regard, TIGAR has been reported to differentially modulate multiple different pathophysiologic outcomes. For example, in the brain TIGAR expression was reported to protect against ischemic/reperfusion injury (Li, Li et al. 2021), whereas in the heart TIGAR deficiency protected myocardial infarction (Hoshino, Matoba et al. 2012) and in pressure overloaded hearts (Okawa, Hoshino et al. 2019). In cancer models, TIGAR plays a complex role on the formation and progression of different types of cancer via suppressing aerobic glycolysis and controlling of reactive oxygen species (ROS) production (Tang, Chen et al. 2021). Recently, it has been reported that TIGAR can both enhance the development of premalignances and suppress the metastasis of cancer invasion by the way of inhibition of ROS production (Cheung, DeNicola et al. 2020).

As carbohydrate metabolism and glycolysis are essential normal physiologic processes in all cells and tissues, we have examined the metabolic phenotype of whole body and tissue specific TIGAR knockout mice. Surprisingly, we have observed that TIGAR deficiency in cholinergic neurons of mice results in a marked protection against hypothermia following an acute cold challenge. At a molecular level, this results from enhanced acetylcholine levels and increased cholinergic signaling at the neuromuscular junction driving skeletal muscle shivering induced thermogenesis.

## Results

Previously we reported that the TIGAR can modulate NF-kB signaling through a direct binding interaction and inhibition of the E3 ligase activity of the linear ubiquitin binding assembly complex, LUBAC in cultured cells (Tang, Kwon et al. 2018). To examine the role of TIGAR in vivo, we initially generated whole body TIGAR knockout mice (TKO) and as expected these mice display a mild increase in adipose tissue inflammation following a high fat diet challenge (data not shown). However, during our phenotypic characterization we observed that the TKO male and female mice both display a remarkable resistance to cold induced-hypothermia. As shown in Figure 1, both male (panel A) and female (panel B) control mice when shifted from room temperature to a 4°C environment display the typical 4-5°C decline in core body temperature during the 1 h acute exposure period. In contrast, the TKO male and female mice only display less than a 1°C decline in core body temperature under the identical conditions. The level of UCP1 mRNA (Figure S1A) and protein (Figure S1B) were not significantly different in interscapular brown adipose tissue (iBAT) at room temperature. UCP1 mRNA did increase approximately 2-fold following 1 h cold exposure but this was not significantly different between the control and TKO mice. Although a very small amount of UCP1 mRNA was detected in inguinal adipose tissue (iWAT) of control mice at room temperature, it was lower in the TKO mice (Figure S1C). Moreover, following 4°C exposure for 1 h there was no significant increase in UCP1 mRNA (data not shown) but by 2 h there was a small measurable increase in UCP1 indicative of beige adipocyte induction. However, the induction of UCP1 was significantly reduced in the TKO mice compared to the control mice, probably due to decrease sensitivity of cold induced sympathetic activation (see Figure 7). As expected, since UCP1 protein levels lag mRNA expression there was no detectable UCP1 protein in iWAT from either control or TKO mice maintained at room temperature or cold exposed for 2 h (data not shown).

**Figure 1:**
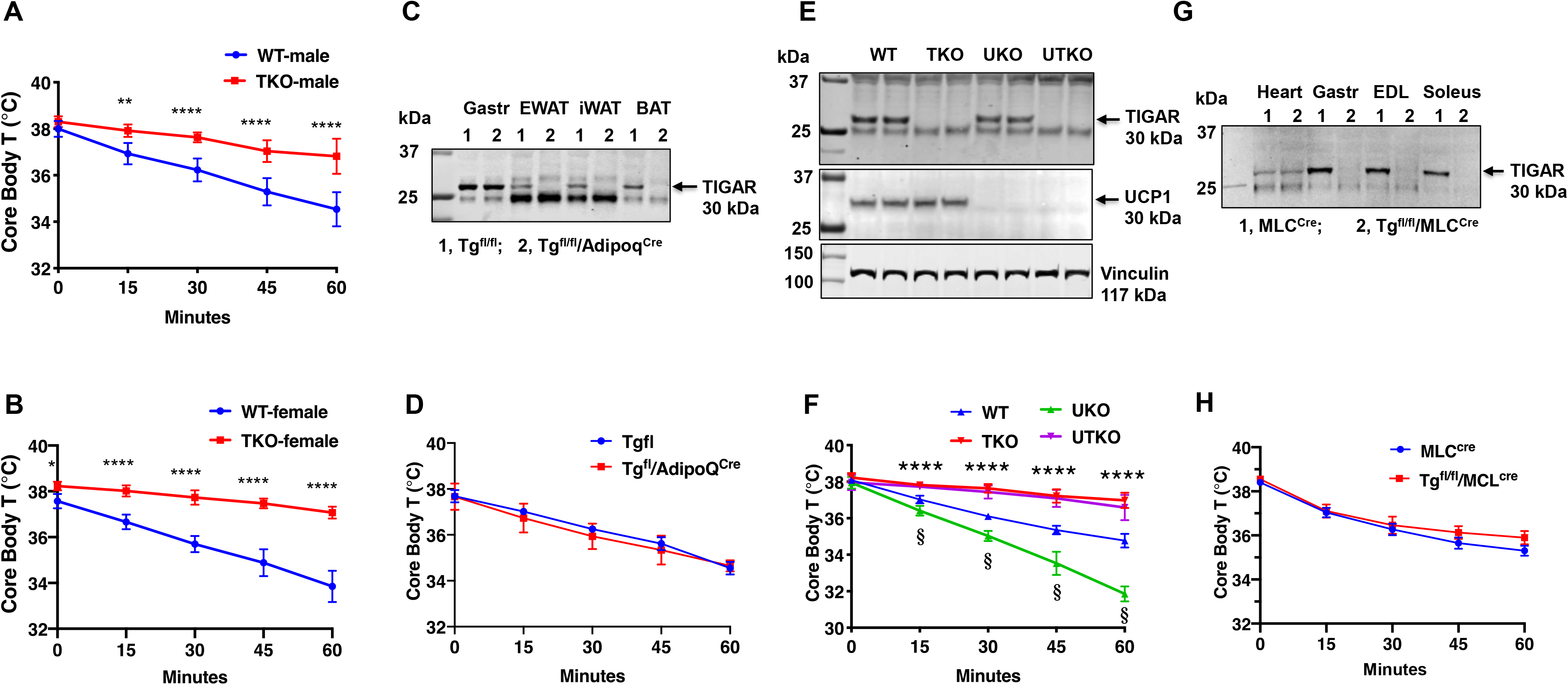
TIGAR deficiency mice are resistant to hypothermia induced by an acute cold challenge. A, Male mice (WT n=10, TKO n=6) and B, Female mice (WT n=6, TKO n=7) core body temperatures were measured every fifteen-minutes starting at ambient laboratory temperature (0 min) and following placement at 4°C. C, Representative TIGAR immunoblot of gastrocnemius muscle (Gastroc), epididymal white adipose tissue (EWAT), subquetaneous inguinal white adipose tissue (iWAT) and interscapular brown adipose tissue (BAT) from control TIGAR flox/flox (Tg^fl/fl^) and adipocyte specific knockout (Tg^fl/fl^/Adipoq^cre^) mice. D, Tg^fl/fl^ and Tg^fl/fl^/AdipoQ^cre^ male mice (n=6) core body temperatures were measured every fifteen-minute starting at ambient laboratory temperature (0 min) and following placement at 4°C. E, Representative UCP1 and TIGAR immunoblots of brown adipose tissue from wild type (WT); whole body TIGAR knockout (TKO), UCP1 knockout (UKO), and UCP1 and TIGAR double knockout (UTKO) mice from two independent genotypes each. F, WT (n=6), TKO (n=6), UKO (n=7), and UTKO (n=8) mice core body temperatures were measured every fifteen-minute starting at ambient laboratory temperature (0 min) and following placement at 4°C. G, Representative TIGAR immunoblot of heart, gastrocnemius muscle (Gastroc), extensor digitorum digitorum longus (EDL) and soleus muscle from control (MLC^cre^) and skeletal muscle specific TIGAR knockout (Tg^fl/fl^/MLC^cre^) mice. H, MLC^cre^ and Tg^fl/fl^/MLC^cre^ male mice (n=6) core body temperatures were measured every fifteen-minute starting at ambient laboratory temperature (0 min) and following placement at 4°C. Statistical analyses was performed using Prism8 and data presented is the mean ± S.D. with two-way ANOVA multiple comparisons. *p* < 0.05, *; *p* < 0.01, **; p<0.001, ***; p<0.0001, ****; p<0.001, ^§^.

As the TKO mice are whole body knockouts, the temperature phenotype could result from an effect on a single or combinatorial number of cell types. We first generated TIGAR^fl/fl^ mice that were crossed with Adiponectin^Cre^ mice to generate adipocyte-specific TIGAR knockout mice (Figure 1C). These mice had the identical temperature sensitivity as the control TIGAR^fl/fl^ mice both displaying an approximate 4°C decline in core body temperature following exposure to 4°C for 1 h (Figure 1D). To further rule out BAT as a contributor to the enhanced cold protection in the TKO mice, we crossed the cold sensitive UCP1 deficient (Enerbäck, Jacobsson et al. 1997) with the TKO mice (both strains with C57BL/6J background) to produce wild type (WT), TKO, UKO, and UCP1^-/-^/TIGAR^-/-^ double (UTKO) mice (Figure 1E). As expected, UKO mice were highly cold sensitive compared to their wild type littermates following a 1 h acute cold exposure (Figure 1F). Interestingly, the UTKO mice displayed a strong cold resistance, that was not significantly different from the TKO littermates. These data strongly suggest that thermogenic adipose tissue is not responsible for the resistance to hypothermia in the TKO mice.

Since skeletal muscle plays a major role in heat production, we next generated skeletal muscle-specific TIGAR knockout mice by crossing the TIGAR^fl/fl^ mice with the skeletal muscle-specific myosin light polypeptide 1-Cre (MLC^cre^) mice (Figure 1G). TIGAR is highly expressed in skeletal muscle, with the highest protein levels in depots containing glycolytic fibers (white gastrocnemius and extensor digitorum longus (EDL) muscles). The MLC^Cre^-driven TIGAR deletion resulted in efficient loss of TIGAR protein in all skeletal muscles examined with no effect on the heart (Figure 1G). Similar to the adipocyte-specific TIGAR knockout mice, skeletal muscle TIGAR deficient mice had no significant resistance to acute cold exposure (Figure 1H). Moreover, we did not find any significant change in skeletal muscle mRNA expression of SERCA1 or SERCA2 in control or TKO mice maintained at room temperature or shifted to 4°C for 1 h (Figure S1D and S1E). Similarly, there was no change in myoregulin (MRLN) (Figure S1F). Although there was a small apparent increase in sarcolipin (SLN) mRNA in the control wild type mice shifted to 4°C there was no statistical difference with the TKO mice (Figure 1G). Together these data are consistent with the absence of a cell autonomous effect in either thermogenic adipocytes or in skeletal muscle.

To unravel these unexcepted findings, we next undertook a pharmacological approach to address the ability of the TKO mice to resist hypothermia. Treatment of control and TKO mice with the selective SERCA inhibitor cyclopiazonic acid (CPA) (Wang, Myles et al. 2021) reduced core body temperature at room temperature (Figure 2A). As expected, CPA treatment resulted in a faster rate of decline and to a lower extent when the mice were exposure to 4°C (Figure 2B). However, there was no significant difference in the response to CPA between the control and TKO mice. The CPA induced reduction in body temperature directly correlated with observable skeletal muscle paralysis (data not shown), consistent with skeletal muscle contraction activity as an important component in thermal regulation (Blondin and Haman 2018).

**Figure 2:**
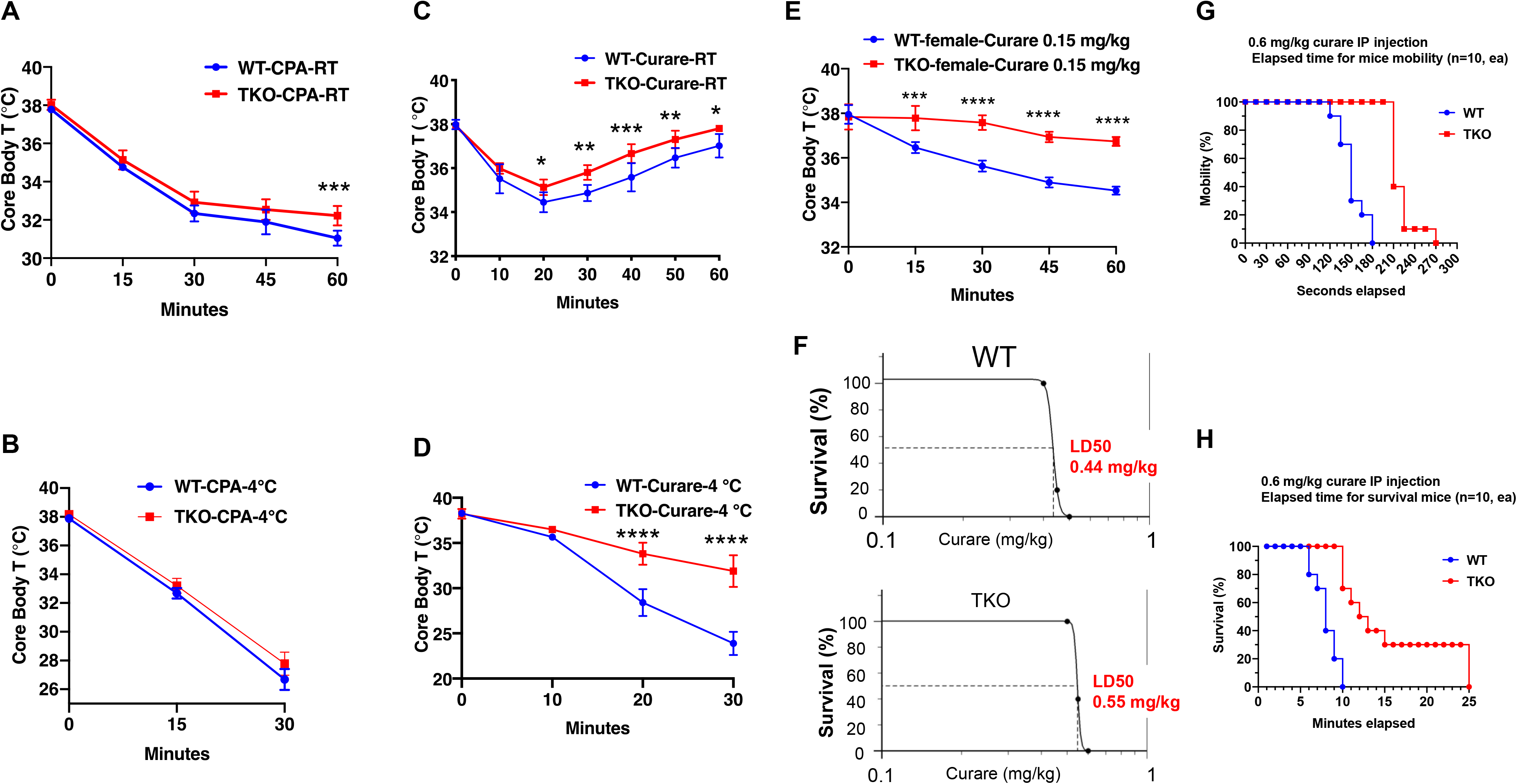
TIGAR knockout mice are resistant to tubocurare but not cyclopiazonic acid. A, Wildtype (WT) and whole body TIGAR knockout (TKO) male mice (n=6) were intraperitoneal injected with cyclopiazonic acid (CPA,10 mg/kg body weight) and then core body temperature was measured every fifteen minutes at room temperature. B, WT and TKO male mice (n=6) were intraperitoneal injected with cyclopiazonic acid (CPA,10 mg/kg body weight) at room temperature and then shifted to 4°C for 30 minutes, with core body temperature measured every fifteen minutes. C, WT and TKO male mice (n=6) were intraperitoneal injected with tubocurare (0.4 mg/kg body weight) and then core body temperature was measured every ten minutes at room temperature. D, WT and TKO male mice (n=6) were intraperitoneal injected with tubocurare (0.4 mg/kg body weight at room temperature and then ten minutes later shifted to 4°C for 30 minutes, with core body temperature measured every ten minutes. E, WT and TKO female mice (n=6) were intraperitoneal injected with tubocurare (0.15 mg/kg body weight) and then core body temperature was measured every fifteen minutes at room temperature. F, WT and TKO male mice (n=10) were intraperitoneal injected with a different tubocurare doses and the LD_50_ of curare was calculated using an online software LD_50_ Calculator, AAT Bioquest, Inc., Sunnyvale, CA. G, WT and TKO male mice (n=10) were intraperitoneal injected with a lethal tubocurare dose (0.6 mg/kg body weight) and the number of mice undergoing complete paralysis was plotted as a function of time in seconds. H, The time to death (absence of respiration) of the same mice was plotted as a function of time in minutes. Statistical analyses were performed using Prism8 and data presented is the mean ± S.D. with two-way ANOVA multiple comparisons. *p* < 0.05, *; *p* < 0.01, **; p<0.001, ***; p<0.0001, ****.

Since TIGAR deficiency in neither adipocytes nor skeletal muscle recapitulated the protection of acute hypothermia but that paralysis results in a rapid reduction in core body temperature, we next assessed whether the cold resistance of the TKO mice was due to shivering thermogenesis. To address this, we next treated mice with the highly specific skeletal muscle nicotinic acetylcholine receptor competitive antagonist tubocurare (Bowman 2006). Control male mice treated with 0.4 mg/kg tubocurare became relatively inactive with essentially no detectable locomotor activity at room temperature 10 min post injection that lasted for another 20-30 min (data not shown). Over this time period there was a concomitant decline in core body temperature that began to recover as the effects of tubocurare began to wear off (Figure 2C). Tubocurare treatment of the TKO male mice at room temperature also displayed a similar decline and recovery of body temperature but that remained somewhat higher than the control mice (Figure 2C). However, shifting the male mice to 4°C following tubocurare injection, resulted in a rapid decline in body temperature such that by 30 min the control mice body temperature was reduced to less than 25°C (Figure 2D). In contrast, the tubocurare injected TKO male mice only display a reduction in body temperature to approximately 34°C under the identical conditions. Moreover, while the tubocurare treated control mice placed at 4°C remained completely immobile, while the TKO mice display apparent normal locomotor activity (Figure S2, movie). Although female mice are more sensitive to tubocurare than male mice (Maurya, Periasamy et al. 2013), TKO female mice were also resistant to the effects of tubocurare on locomotor activity (data not shown) and acute cold exposure (Figure 2E).

The tubocurare dose-response for male mice is shown in Figure 2F, with the control wild type mice have an LD50 of 0.44 mg/kg whereas the TKO mice have an LD50 of 0.55 mg/kg (Figure 2F). Injection of a lethal tubocurare dose (0.60 mg/kg) initially resulted in 50% if the control mice displaying a loss of voluntary locomotor activity by 150 sec but this required 240 sec in the TKO mice (Figure 2G). At ten min 100% of the control mice were dead whereas it took nearly 25 min for 100% of the TKO mice to die (Figure 2H). The tubocurare resistance of the TKO mice was not due to changes in expression of the nicotinic acetylcholine receptor subunits with no significant differences in the skeletal muscle expression of the α1, β1 or γ subunits (Figure S3A, 3B, and 3C) and with a small reduction in the δ and ε subunits (Figure S3D and S3E) in the TKO mice. These data suggest that the protection against hypothermia was due to cholinergic signaling to the skeletal muscle.

Consistent with increased skeletal muscle energetic demand, ATP levels were significantly decreased with an apparent increase in ADP and AMP in the TKO mice after 1 h at 4°C (Figure 3A). Stable isotope metabolic flux assessment with H_2_O^18^ demonstrated increased incorporation of orthophosphate into ATP, ADP and AMP in the TKO mice at 4°C, that was not significantly different at room temperature (Figure 3B). Plasma creatine was also increased 2-fold in the TKO at 4°C (Figure 3C), reflecting the increased phosphocreatine turnover (CP M4 phosphate fragment m/z 83, Figure 3D). The appearance of creatine in the plasma is likely due, in part, to the increase rate of shivering that is known to induce skeletal muscle leakage (Baird, Graham et al. 2012). The mechanics of energetic supply at 4°C was further examined by using a widely targeted LC/MS/MS protocol (Hoshino, Matoba et al. 2012). Key pathways found to be significant were pentose phosphate pathway, purine nucleotide cycle, and amino acid utilization pathways (Figure S4A). The pentose phosphate pathway cycle fills the purine nucleotide cycle that acts in concert with the malate-aspartate shuttle and the TCA cycle to preserve cellular energetics (Hatazawa, Senoo et al. 2015). In particular glycine/serine, histidine and methionine metabolism also drives the input of the tricarboxylic acid cycle intermediates serving as anapleurotic inputs (Figure S4B). Together, these interactions serve to recycle AMP generated by increased skeletal muscle energy demand.

**Figure 3:**
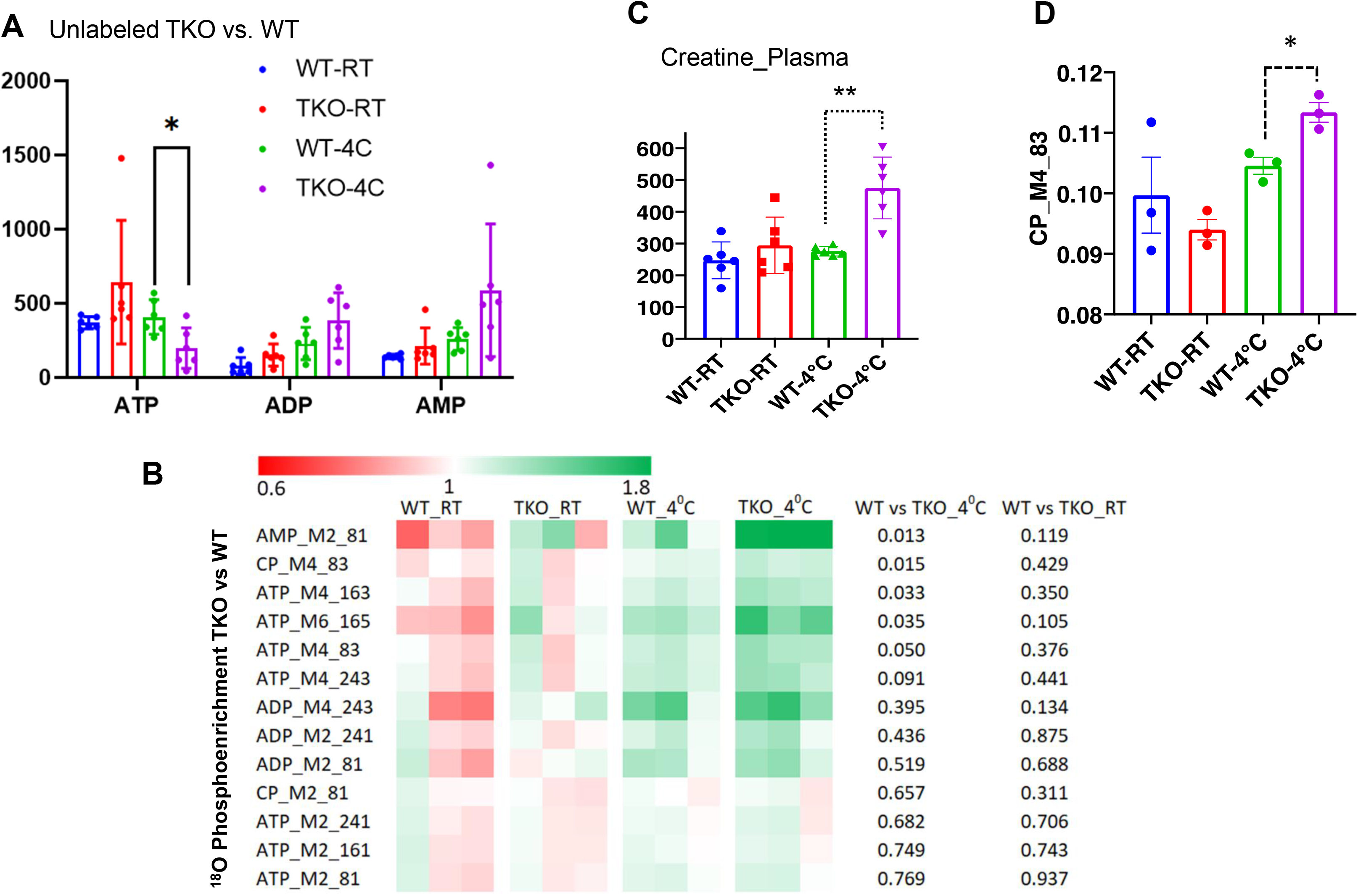
TIGAR deficient mice display increased ATP turnover in skeletal muscle. A, Wildtype (WT) and TIGAR knockout (TKO) male mice (n=6) were maintained at room temperature (RT) or shifted to 4°C for 1 hour. Quadriceps white skeletal muscle was isolated and extracts assayed for ATP, ADP and AMP levels as described in Method and Materials. B, Wildtype (WT) and TIGAR knockout (TKO) male mice (n= 3-4 per condition) were at room temperature or shifted to 4°C for 1 h and then given an oral gavage of 0.3 ml pure H_2_O^18^ for 10 min at 4°C and an IP injection of 1 ml pure H_2_O^18^ for another 10 min at 4°C. The quadricep muscles were freeze-clamped with liquid N_2_ and extracts prepared for mass spectroscopic analyses. The heatmap shows the O^18^ enrichment fraction of ATP, ADP, AMP, and creatine-phosphate. C, Wildtype (WT) and TIGAR knockout (TKO) male mice (n=6) were maintained at room temperature or shifted to 4°C for 1 hour and plasma creatine levels were determined by metabolomics analyses. D, The CP_M4_83 data was generated from the same experimental setting as B, indicating the increase of phosphocreatine turnover in TKO compared to that in WT at 4°C. Statistical analyses were performed using Prism8 and data presented is the mean ± S.D. with two-way ANOVA multiple comparisons. *p* < 0.05, *; *p* < 0.01, **.

Mitochondria are a primary source of both ATP production and heat generation during muscle contraction. To determine whether mitochondrial oxidative phosphorylation (OXPHOS) efficiency and/or capacity are altered in TKO mice, permeabilized skeletal muscle fiber bundles were prepared from red portions of the gastrocnemius muscle from control and TKO mice immediately after 1 h cold exposure. ADP-stimulated respiratory kinetics (i.e., Km and Vmax) were virtually identical between wild-type and TKO muscle during respiration supported by carbohydrate (Figure S5A and S5B) and/or lipid-based substrates (data not shown). Rates of ATP production (*J*ATP) and oxygen consumption (*J*O_2_) measured simultaneously under clamped submaximal and maximal ADP-demand states were also similar in muscle from control and TKO mice (Figure S5C and S5D), yielding similar ATP/O efficiency ratios (Figure S5E). Finally, in vitro studies of the extensor digitorum longus muscle showed no differences in force-frequency, peak specific tension, or time to one-half relaxation between control and TKO mice (Figure S5F and S5G). Together these data indicate that the skeletal muscle contraction induced thermogenesis in TKO mice is not due to any intrinsic change in skeletal muscle mitochondrial efficiency of contractile function and therefore must result from a change in signaling that in turn drives an enhanced skeletal muscle response.

Since the TKO mice are resistant to tubocurare and the major driver of skeletal muscle contraction is cholingeric signaling, we next generated cholinergic neuron specific TIGAR knockout (chTKO) mice using the choline acetyl-CoA transferase (ChAT)-Cre mice (Rossi, Balthasar et al. 2011). We confirmed that ChAT^Cre^ resulted in the loss of TIGAR in cholinergic neurons by immunofluorescence co-localization of TIGAR with the vesicular choline transporter (VChAT) (Figure 4A). As the superior cervical ganglion (SCG) is composed of approximately 80% cholinergic neurons (Wang, Nelson et al. 1990, Morales, Holmberg et al. 1995, Juranek and Wojtkiewicz 2015), immunoblotting also confirmed the efficient loss of TIGAR protein (Figure 4B). Similar to the TKO mice, the chTKO mice were also resistant to acute cold exposure compared to the control ChAT^Cre^ mice (Figure 4C). Treatment with the SERCA inhibitor CPA, at room temperature Figure 4D) or at 4°C (Figure 4E), also resulted in paralysis and a decline in core body temperature that was not significantly different between the ChAT^Cre^ and chTKO mice, although the reduction in core body temperature was much greater at 4°C than at room temperature. The chTKO mice were also resistant to tubocurare at room temperature with both less of a core body temperature drop and a faster recovery (Figure 4F). In agreement with the TKO mice (Figure 2D), the resistance to cold induced hypothermia by acute 4°C challenge was very dramatic in the chTKO mice compared to control mice (Figure 4G). Similarly, the chTKO mice were highly resistance to the paralytic actions of tubocurare (Figure S6, movie). Moreover, immunoblotting of the SCG for the neuron activation marker c-Fos, further demonstrated that control mice displayed a small increase in c-Fos protein levels when cold challenged that was further increased in the chTKO mice (Figure 4H). These data demonstrate the cholinergic neuron TIGAR knockout mice recapitulate the cold resistance and pharmacological characteristics of the whole body TIGAR knockout mice, consistent with these mice displaying increased cholinergic tone.

**Figure 4:**
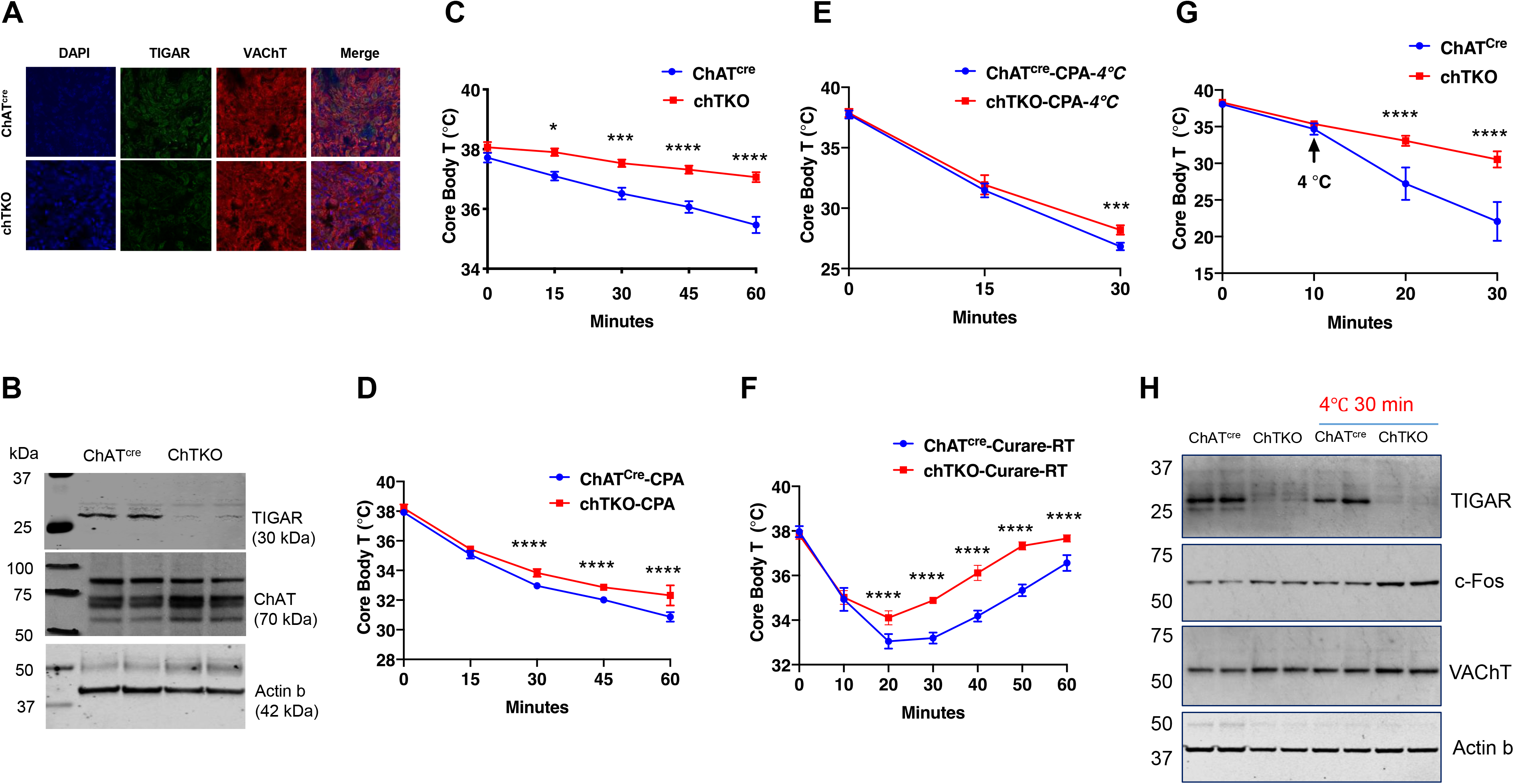
Cholinergic neuron specific TIGAR knockout mice recapitulate the protection against hypothermia of the whole-body TIGAR deficient mice. A, Representative immunofluorescent images showing TIGAR, vesicular acetylcholine transporter (VAChT), and nuclei (DAPI) in the superior cervical ganglions (SVG) from control ChAT^cre^ and Tg^fl/fl^/ChAT^cre^ (chTKO) male mice. B, Representative TIGAR, choline acetyltransferase (ChAT) and actin protein immunoblots of the SVG from two independent control (ChAT^cre^) and chTKO male mice. C, Male ChAT^cre^ and chTKO mice (n=6) core body temperatures were measured every fifteen-minutes starting at ambient laboratory temperature (0 min) and following placement at 4°C. D, Male ChAT^cre^ and chTKO mice (n=6) were intraperitoneal injected with cyclopiazonic acid (CPA,10 mg/kg body weight) at room temperature and core body temperature measured every fifteen minutes. E, ChAT^cre^ and chTKO male mice n=6) were intraperitoneal injected with cyclopiazonic acid (CPA,10 mg/kg body weight) at room temperature, shifted to 4oC and core body temperature measured every fifteen-minute. F, Male ChAT^cre^ and chTKO mice (n=6) were intraperitoneal injected with tubocurare (0.4 mg/kg body weight) and then core body temperature was measured every fifteen minutes at room temperature. G, Male ChAT^cre^ and chTKO mice (n=6) were intraperitoneal injected with tubocurare (0.4 mg/kg body weight at room temperature and ten minutes later shifted to 4°C for 30 minutes. Core body temperature was measured every ten minutes. H, Representative TIGAR, c-Fos, VAChT and actin protein immunoblots of the SVG from two independent control (ChAT^cre^) and chTKO male mice at room temperature or shifted to 4°C for 30 min. Statistical analyses were performed using Prism8 and data presented is the mean ± S.D. with two-way ANOVA multiple comparison. *p* < 0.05, *; p<0.001, ***; p<0.0001, ****.

If increased cholinergic input is responsible for the increased energy demand and heat production by skeletal muscle, then the skeletal muscle metabolic characteristics of the TKO mice should be recapitulated in the chTKO mice. As shown in Figure 5, widely targeted LC/MS/MS in the gastrocnemius muscle of the chTKO mice identified the same three key pathways: pentose phosphate pathway, purine nucleotide cycle, and amino acid utilization pathways that display a greater induction at 4°C. Together, these data are consistent with increased cholinergic signaling to skeletal muscle as responsible for increased skeletal muscle thermogenesis following cold exposure.

**Figure 5:**
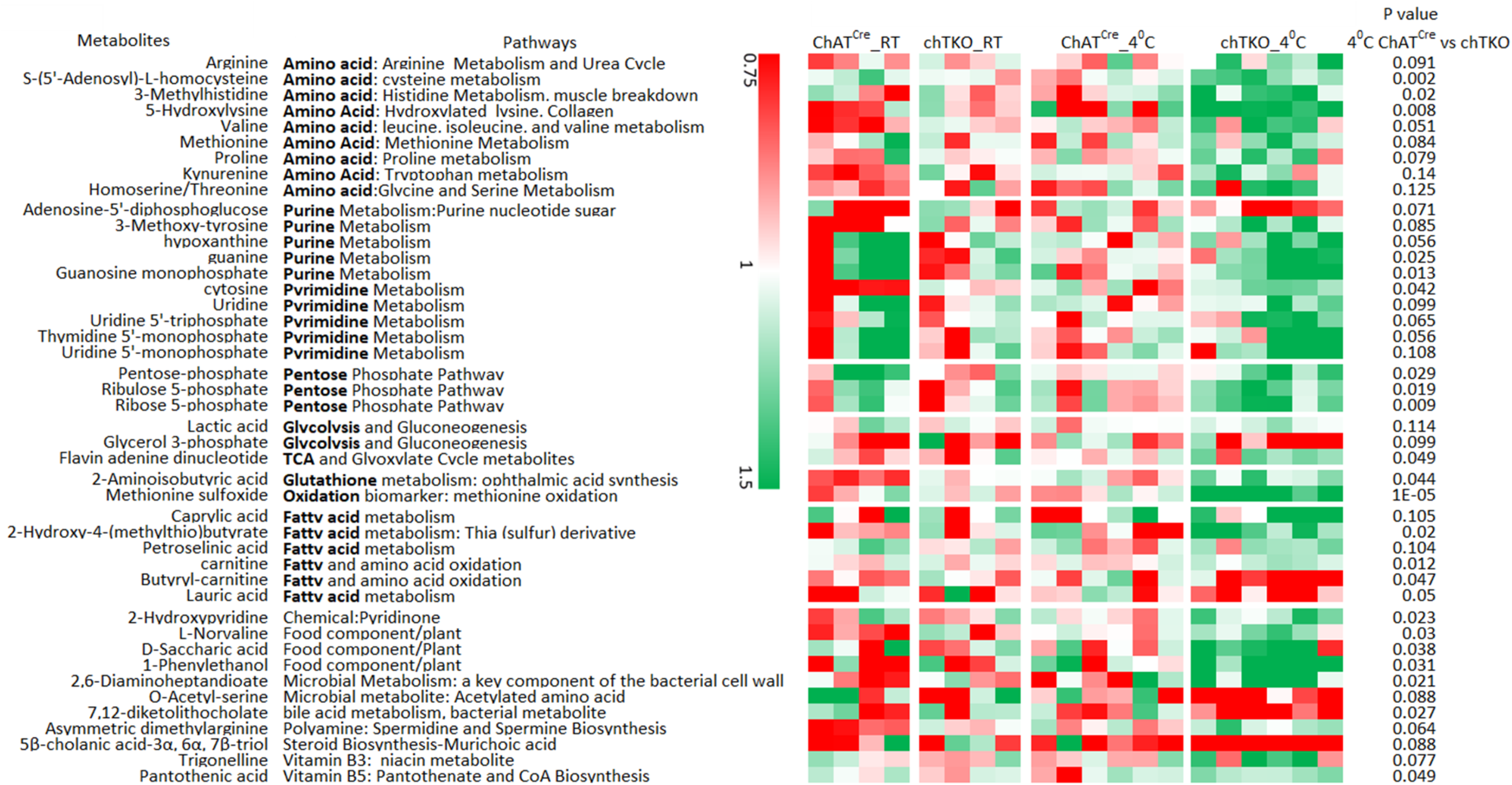
Skeletal muscle of chTKO mice at 4°C display increased pentose phosphate pathway, purine nucleotide cycle, and amino acid utilization pathways. Control (ChAT^cre^) and chTKO male mice (n=4-6) were maintained at room temperature (RT) or shifted to 4°C for 1 hour. Quadriceps white skeletal muscle was isolated and extracts were subjected to widely targeted (MRM) small metabolite profiling using an ABSciex 6500+ QTRAP with ACE PFP and MERCK Zic-pHILIC columns as described in Method and Materials. The heatmap shows the metabolites/pathways differentially identified with corresponding p values.

To address the basis of the apparent increase in cholinergic signaling, we first asked whether there was a change in synaptic density in the neuromuscular junction. The acetylcholine receptor aggregates in the synaptic cleft and can be specifically labeled with α-bungarotoxin (Pratt, Iyer et al. 2018), as shown for whole extensor digitorum longus (EDL) muscle (Figure 6A). Quantification revealed a large degree of heterogeneity in the number of clustered acetylcholine receptors per muscle (Figure 6B). Nevertheless, there was a clear statistical increase in the TIGAR knockout mice compared to control mice. Consistent with the increase in synaptic junctions, the total amount of acetylcholine was also increased in skeletal muscle extracts from the TKO and chTKO mice compared to their respective control mice at both room temperature and after cold exposure (Figure 6C and 6D).

**Figure 6:**
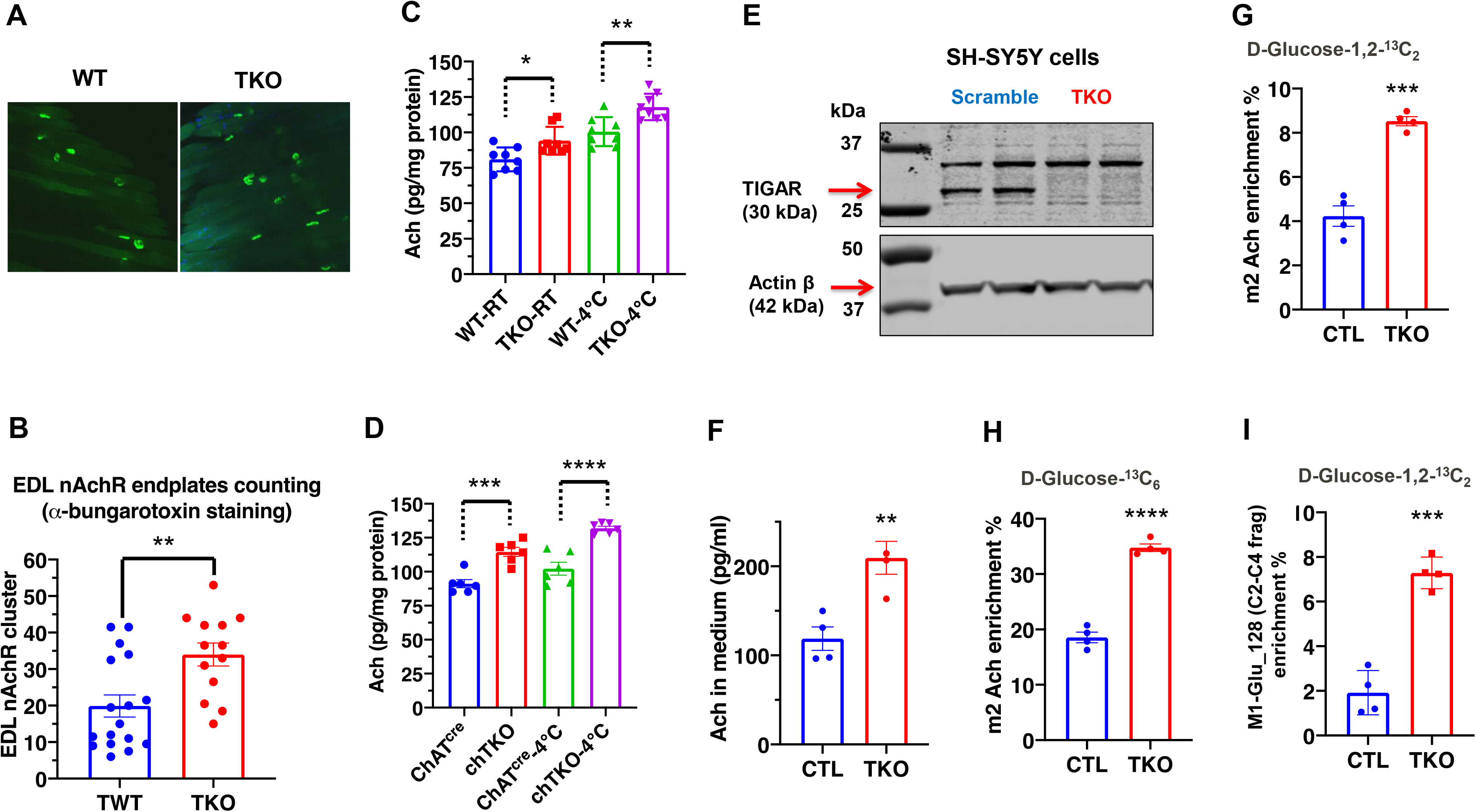
TIGAR deficiency increases acetylcholine biosynthesis and acetylcholine receptor clustering at the neuromuscular junction. A, Representative images of a-bungarotoxin immunofluorescent labeling of nicotinic acetylcholine receptor clusters in EDL muscle from wildtype (WT) and TKO mice. B, Quantification of the number of nicotinic acetylcholine receptor clusters following fifteen minutes exposure at 4°C (WT n=17, TKO n=13). These data represent the average of over six mice in each group of mean ± S.D. (Unpaired t test, two-tailed, p=0.0035, **). C, Acetylcholine levels in gastrocnemius muscle of WT and TKO male (n=7) mice at room temperature or following 1 hour at 4°C. D, Acetylcholine levels in gastrocnemius muscle of ChAT^cre^ and chTKO male (n=6) mice at room temperature or following 1 hour at 4°C. E, Representative immunoblots of TIGAR and actin proteins from two scrambled sgRNA and two TIGAR sgRNA knockout Sh-SY5Y cell lines. F, Acetylcholine concentrations in the medium of scrambled and TKO SH-SH5Y neuroblastoma cells. G, m2 Acetylcholine enrichments in cells labeled with D-glucose-1,2-^13^C_2._ H, m2 Acetylcholine enrichments in cells labeled with U-^13^C_6_ D-glucose-. I, m1 Glutamate (m/z 128, C2-C4 fragment) enrichment in the medium of scrambled and TKO SH-SY5Y human neuroblastoma cells labeled with D-glucose-[1,2]-^13^C_2_. Statistical analyses were performed using Prism8 and data presented is the mean ± S.D. with two-way ANOVA multiple comparison. *p* < 0.05, *; p<0.001, ***; p<0.0001, ****.

As it is not possible to determine the rate of acetylcholine synthesis in vivo, we took advantage of SH-SY5Y cells that have characteristics of cholinergic neurons (Kovalevich and Langford 2013). Using sgRNA and CAS9, we effectively knocked out TIGAR in several clonal SH-SY5Y cells, two representative control and sgRNA knockout cells are shown in Figure 6E. The amount of acetylcholine secreted into the medium over a 24 h period was increased in the TIGAR knockout SH-SY5Y cells compared to controls (Figure 6F). Labeling with [1,2]-^13^C glucose demonstrate an increase in the m2-labeled acetylcholine levels in the TIGAR knockout SH-SY5Y cells (Figure 6G). Similarly, labeling with U-^13^C glucose also demonstrated an increase in m2-labeled acetylcholine in the TIGAR knockout SH-SY5Y cells (Figue 6H). Moreover, there was a 3-fold increase in the enrichment of the C2-C4 fragment, m1-glutamate m/z 129), indicating that the acetyl group in acetylcholine originated from glycolysis via pyruvate dehydrogenase, PDH (Figure 6I, Figure S7). This is because the m2 acetyl group formed via PDH generates ^13^C atoms at positions 4 and 5 of glutamate, wheras the m2 ^13^C glutamate at position 2 and 3 is derived from pyruvate carboxylase (PC) generated oxaloacetate (Madhu, Boneski et al. 2020) schematically shown in Figure S7A. Analyses of the m/z 128 C2-C4 glutamate fragment distinguishes these isotopomers with the enrichment of m1 glutamate (129 fragment) approximately 5 times more abundant than the m2 glutamate (130 fragment) in the TIGAR knockout cells (Figur S7B). Together, these data demonstrate that TIGAR deficiency in SH-SY5Y results from an increase in acetyl-CoA levels consistent with increased glycolysis and direct conversion of pyruvate to acetyl-CoA by pyruvate dehydrogenase, as opposed to pyruvate carboxylase and subsequent entry into the tricarboxylic acid cycle.

A generalized increased cholinergic tone would be expected to reduce blood pressure and heart rate but with increased skeletal muscle shivering. To assess these physiologic parameters, we used implanted telemetry probes to simultaneously determine body temperature, mean arterial pressure (MAP), heart rate (BPM), and skeletal muscle electromyography in cold exposed ChAT^cre^ and chTKO mice (Figure 7). Consistent with the rectal temperature measurements previously observed, the telemetry probe also demonstrated that the chTKO are more resistant to cold induced hypothermia than ChAT^cre^ mice (Figure 7A). At room temperature, there was no significant difference in blood pressure, but following cold exposure the blood pressure of the control mice increased whereas the blood pressure of the chTKO mice decreased such that by 60 min the chTKO mice had a blood pressure 25 mmHg less than the control mice (Figure 7B). Similar, the heart rate was not statistically different at room temperature but increased substantially more in the control mice than in the chTKO mice following cold exposure (Figure 7C). As these data strongly suggest increase cholinergic signaling to enhance skeletal muscle contraction, we utilized electromyography (EMG) to directly determine skeletal muscle activity. A representative image of EMG tracers and root mean square (RMS) derivation are shown for ChAT^cre^ and chTKO mice when shifted to 4°C for 30 min (Figure 7D). Quantification of these types of data as the integral of RMS during the time course of cold exposure, demonstrated that at room temperature and following 10 min of cold exposure there is no significant difference in skeletal muscle contraction activity between ChAT^cre^ and chTKO mice (Figure 7E). However, the contraction activity of the control ChAT^cre^ mice plateau at this level, whereas the chTKO mice shivering activity further increases to nearly twice the level of the control mice. Together, the relative changes in cold induced blood pressure, heart rate, and skeletal muscle shivering are fully consistent with the chTKO mice displaying a generalized increase in cholinergic signaling following cold exposure.

**Figure 7:**
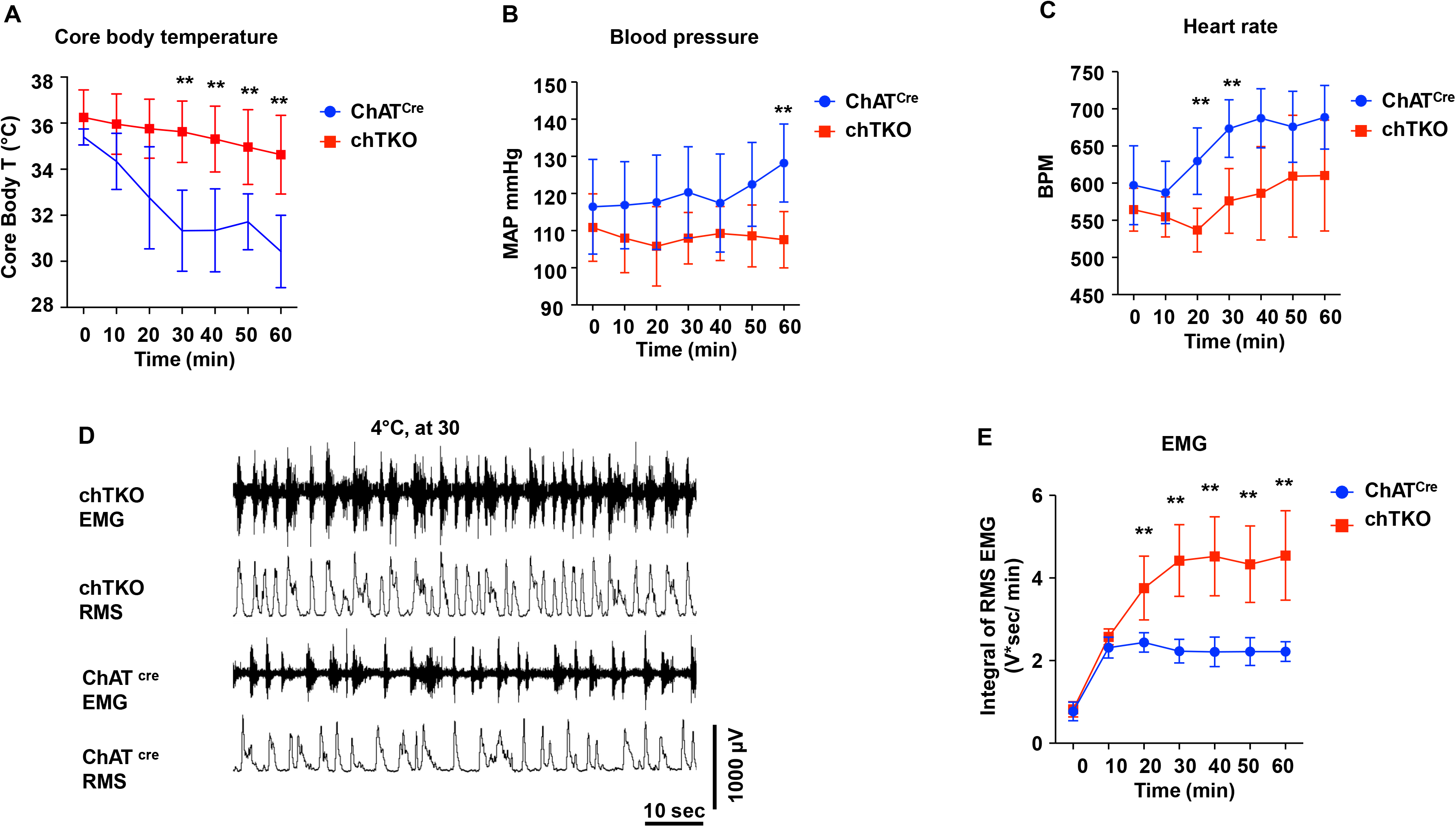
ChTKO mice display enhanced cold stimulated skeletal muscle shivering activity and increased cholinergic tone. A, Male mice core body temperatures (ChAT^cre^ n=7, TKO n=10), B, blood pressure (mmHg) (ChAT^cre^ n=7, chTKO n=10), C, heart rate (beats per minute, BPM) (ChAT^cre^ n=7, chTKO n=6), D, Representative electromyography traces (EMG) and root mean squares (RMS) for ChAT^cre^ and chTKO male mice when shifted to 4°C for 30 min, and E, neck electromyogram (EMG, (ChAT^cre^ n=5, chTKO n=6)) were measured continuously starting at ambient laboratory temperature (0 min) and for 60 min following placement at 4°C. Statistical analyses were performed using Prism 9 and data presented are the means ± S.D. with two-way ANOVA multiple comparisons. *p* < 0.01*.

## Discussion

Currently, there is a substantial effort to understand and develop approaches to increase energy expenditure through the development/activation of brown/beige adipose tissue thermogenesis. A fraction of adult humans can respond to cold stress by inducing the expression of thermogenic adipose tissue under relatively long-term cold acclimation in adult humans (McNeill, Suchacki et al. 2021). However, it remains somewhat controversial if this inducible thermogenic human brown adipose tissue is the rodent equivalent of beige or brown adipocytes (Cannon, de Jong et al. 2020, Samuelson and Vidal-Puig 2020, Virtanen and Nuutila 2021). In addition, whether the mass of the human inducible thermogenic adipocytes is sufficient to significantly contribute to cold responsive thermogenesis for core temperature maintenance or weight loss has also been questioned (Blondin, Tingelstad et al. 2014, Virtanen and Nuutila 2021). In contrast, skeletal muscle accounts for approximately 40% of healthy body mass, is the dominant factor for basal metabolic rate and the major driver of energy expenditure during physical exercise (Haman, Blondin et al. 2010, Blondin, Tingelstad et al. 2014). In rodents, non-shivering thermogenesis (NST) is driven by UCP1 dependent mitochondrial uncoupling in both brown and beige adipocytes, by the creatine phosphorylation/dephosphorylation cycle in beige adipocytes (Cohen and Spiegelman 2015, Kazak, Chouchani et al. 2015, Jung, Sanchez-Gurmaches et al. 2019). There are also reports of a skeletal muscle based NST via a sarcolipin-dependent futile cycle of SERCA-dependent calcium transport in the sarcoplasmic reticulum (Bal, Maurya et al. 2012, Maurya, Bal et al. 2015). In humans, maximal NST can increase energy production equivalent to about 2 times above the resting metabolic rate (RMR), while moderate shivering thermogenesis (ST) is 2.5 to 3.5 times RMR and maximal shivering has been measured at about 5 times RMR (Haman, Blondin et al. 2010, Blondin, Tingelstad et al. 2014). In contrast, exercise at ∼50% VO_2_ max generates approximately 15 times RMR (Weber and Haman 2005, Blondin, Tingelstad et al. 2014). While a substantial effort has focused on NST, shivering and physical activity are the dominant forms of involuntary and voluntary heat production during acute cold exposure. Thus, it is essential to develop a deeper understanding of the physiologic integrated role of skeletal muscle with thermogenic adipose tissue to have a complete and detailed molecular understanding of energy balance and its implications for potential interventions to promote and maintain weight loss.

The data presented in this manuscript demonstrate that cholinergic neuron deficiency of TIGAR results in an enhancement of cold-induced skeletal muscle contraction/shivering that is responsible for an increase in heat generation and protection of acute cold induced hypothermia. This response is likely mediated by an increase acetylcholine signaling at the neuromuscular junction, consistent with the parallel induction of tubocurare resistance and increased cholinergic neuron activation. Interestingly, despite the increase in synaptic density and acetylcholine levels mice at room temperature that are mildly cold stress (thermoneutrality for mice is approximately 29-31°C (Ganeshan and Chawla 2017)), TIGAR deficiency does not appear to significantly affect body temperature, skeletal muscle contraction activity or the overall skeletal muscle metabolite profile at room tempature. However, following acute cold challenge there are marked changes in these parameters between wild type and cholinergic TIGAR knockout mice including a metabolic profile indicative of increased ATP turnover. These findings suggest that there is an increase in cholinergic signaling reserve that only becomes apparent when cholinergic signaling demand is high.

In addition to TIGAR as a small carbohydrate phosphatase, previous studies have shown that TIGAR can directly interact with hexokinase 2 and the HOIP subunit of the linear ubiquitin assembly complex (Cheung, Ludwig et al. 2012, Tang, Kwon et al. 2018). Thus, although it is formally possible that TIGAR increases cholinergic signaling due to alterations in protein-protein interactions, we believe it is more likely a result of increased glycolysis due to its phosphatase function, that is occurring in the cholinergic neurons as this would result in an increase in acetyl-CoA levels. The steady-state acetyl-CoA concentrations in neuronal mitochondrial and cytosol are several folds lower than acetyl-CoA Km values for choline acetyltransferase. More specifically, following cholinergic differentiation there is a further decrease in acetyl-CoA levels that results in further a substrate-dependence of acetylcholine biosynthesis (Ronowska, Szutowicz et al. 2018). Thus, the rate-limiting step of acetylcholine synthesis is the relative concentration of acetyl-CoA and since TIGAR deficiency increases the amount of acetyl-CoA through enhanced glycolysis, this leads to increased production of acetylcholine in cholinergic neurons. At present, it is not possible to distinguish the amount of synaptic acetyl-CoA from skeletal muscle acetyl-CoA at the neuromuscular junction, but future studies using spatial mass spectroscopy (MALDI) may be able to resolve this issue.

Finally, it is also important to recognize that myasthenia gravis is a neuromuscular disease caused by autoantibodies against components of the neuromuscular junction and in particular the nicotinic acetylcholine receptor (Gilhus, Tzartos et al. 2019). Current therapeutic strategies are focused on the use of steroids or immune suppressants to block the immune system and drugs to increase the transmission of acetylcholine signaling such as acetylcholine esterase inhibitors (Gilhus, Tzartos et al. 2019). Similarly, central cholinergic neurons undergo severe neurodegeneration in Alzheimer’s disease and agents that enhance cholinergic signaling, particularly cholinesterase inhibitor therapies, provide significant symptomatic improvement in patients with Alzheimer’s disease (Ferreira-Vieira, Guimaraes et al. 2016, Ahmed, Knowles et al. 2019). Interestingly, recent reports show that the neuromuscular junctions are stable in human age-related sarcopenia and in patients with cancer cachexia, which are different with the rodent models in that the sarcopenia is linked with the endplate fragmentation in NMJ (Jones, Harrison et al. 2017, Boehm, Miller et al. 2020). As such, inhibition of TIGAR function may also provide a novel approach to improve skeletal muscle function, memory and dementia through increased cholinergic signaling while limiting the deleterious metabolic consequences secondary to currently available therapies.

## Acknowledgement

We thank Dr. Yu Zhang at the Flow cytometry Core Facility (Albert Einstein College of Medicine) for SH-SY5Y cells FACS sorting. We also thank Dr. Victor Schuster (Albert Einstein College of Medicine) for the gift of UCP1 knockout mice. We thank Ms. Nicole Fernandez (Ph.D. candidate at the Albert Einstein College of Medicine) for her contribution on the production of Graphic Abstract. This study was supported by grants DK033823 and DK020541 from the National Institutes of Health, and a S10 SIG Award for the Sciex 6500+ QTRAP (1S10OD021798).

## Author contributions

Y. T., H. Z., H. W. and Y. Q. designed and conducted experiments, compiled/analyzed and interpreted data, prepared figures, and wrote and edited the manuscript. J. B. P. assisted in experimental design, compiled/analyzed and interpreted data, and wrote and edited the manuscript. L. W., K. A. B., I. B., C., A. S., and CT. L. conducted experiments, compiled/analyzed and interpreted data, prepared figures. P. D. N., G. J. S., and I. J. K. designed and conducted experiments, compiled/analyzed and interpreted data, prepared figures, and wrote and edited the manuscript. J. E. P. was responsible for the overall direction, experimental design, analyses and interpretation of data, writing and editing of the manuscript, and funding acquisition.

## Materials and Methods

### Resource availability

#### Lead contact

Further information and requests for resources and reagents should be directed to and will be fulfilled by the lead contact, Jeffrey E Pessin

#### Materials availability

This study did not generate new unique reagents. SH-SY5Y neuroblastoma both scramled and TKO cell lines are available upon request from the lead contact. Mouse lines generated in this study are available from the lead contact, although there are timeline restrictions to the availability of the mice due to the labor and space limitations of the laboratory staff and the facility of Institution for Animal Studies at Albert Einstein College of Medicine.

#### Data and code availability

The authors are willing to deposit all the data in this study in datatype-specific, Cell Press-recommended repository before the manuscript being accepted for publication.

### Experimental models and subject details

#### Mice models

All studies were performed in accordance with protocols approved by the Einstein Institutional Animal Care and Use Committee. TIGAR whole body knockout (TKO) mice were made as described in our previous paper (Tang, Kwon et al. 2018). To establish conditional knockout mice of *TIGAR* gene, we purchased C57BL/6N-^TIGARtm1b(EUCOMM)Wtsi^/Wtsi mice from Welcome Trust Sanger Institute (Hinxton Cambridge, U.K.). Briefly, loxP sites were generated to delete exon 3 of *TIGAR*, chimeric mice were mated with C57BL/6 mice and germ-line transmission confirmed by PCR and Southern blotting. The progeny crossed sequentially with *FLPe* transgenic mice to remove lacZ and neo genes. The primer pairs TIGARCKO-5FRT-FW: TTGGGATCCCCTAGTTTGTG and TIGARCKO-5FRT-RV: AACTCAGCCTTGAGCCTCTG were used to confirm deletion of the lacZ and neomycin resistance genes. The primer pairs TIGARCKO-3loxp-FW: GAGAAGAGACCCCCTGGAAC and TIGARCKO-3loxp-RV: TTCCGGCCAAACAGACTTAC were used to confirm the *TIGAR*^f/f^ mice. Mice were backcrossed more than ten generations to C57BL/6J mice. The primers for subsequent genotyping for *TIGAR*^f/f^ were listed in the Key Resources Table (KRT) and the mouse tail PCR (touchdown cycling protocol, Jackson Laboratory) results in 538 bp for mutant and 346 bp for wild type mice.

Adipocyte-specific TIGAR knockout (AdipoqTKO) mice were produced by mating TIGAR^f/f^ mice with adiponectin-Cre mice, as previously described (Feng, Amgalan et al. 2018). Skeletal muscle-specific TIGAR knockout mice were produced by mating the *TIGAR*^f/f^ mice with skeletal muscle-specific myosin light polypeptide 1 *Myl1*^cre^ mice purchased from Jackson Laboratory (stock number 024713). Cholinergic neuron-specific TIGAR knockout (chTKO) mice were generated by mating TIGAR^f/f^ mice with ChAT^cre^ mice purchased from Jackson Laboratory (stock number 028861). UCP1 knockout (UKO) mice were a gift from Dr. Victor Schuster (Albert Einstein College of Medicine). UCP1/TIGAR double knockout (UTKO) mice were generated by breeding UCP1 knockout mice with TKO, which resulted in the production of WT, TKO, UKO, and UTKO mice.

#### Mouse husbandry and rectal temperature measurement

Mice were housed in a facility equipped with a 12-h light/dark cycle. Animals were fed a normal chow diet that contains 62.3% (kcal) carbohydrates, 24.5% protein, and 13.1% fat (5053, LabDiet). The mice at 12-16 weeks of age were used for rectal temperature measurement using TH-8 Thermometer (Physitemp Instruments, Clifton, NJ) per manufacturer’s instructions. For cold exposure, the mouse was individually put into a pre-chilled cage with bedding in a cold chamber (4°C). For tissue collection, the mice were killed, and the tissues were collected and snap-frozen in liquid nitrogen and stored in a −80°C freezer. All studies were approved by and performed in compliance with the guidelines of the Albert Einstein College of Medicine Institutional Animal Care and Use Committee.

#### Blood plasma collection

Mice were briefly anesthetized by isoflurane and the blood sample was collected from mouse orbital sinus into microtube with EDTA (Microvette 500 K3E, SARSTEDT, Germany) on ice. The plasma sample was collected by centrifugation (1,000 g, 30 min) and was stored at −80°C.

#### Gastrocnemius and quadricep skeletal muscle samples isolation

Gastrocnemius and quadricep white muscle were collected and snap-frozen in liquid nitrogen and stored at −80 °C. The whole piece of the muscle was ground using a mortar-pestle in liquid nitrogen and the muscle powder (30 mg) was used for metabolomics analysis.

#### Tubocurare and cyclopiazonic acid treatments

Tubocurare (T2379, Sigma-Aldrich) was given as an intraperitoneal injection in a total volume of 100 μl in saline solution reaching at various doses and times indicated in the figure legends and the median lethal dose (LD50) was calculated with LD50 Calculator, a online tool from AAT Bioquest (Sunnyvale, CA). Cyclopiazonic acid (C1530, Sigma-Aldrich) was given as an intraperitoneal injection in a total volume of 100 μl in saline solution reaching a dose of 10 mg/kg body weight for the time indicated. In both cases, the same volume of 100 μl of saline were injected as vehicle controls.

#### TIGAR knockout in human SH-SY5Y neuroblastoma cells

Human SH-SY5Y neuroblastoma cell line was purchased from ATCC (ATCC^®^ CRL-2266). The SH-SY5Y cells were cultured in Dulbecco’s modified Eagle’s medium (DMEM) supplemented with 10% fetal bovine serum and 1× penicillin-streptomycin. Cell lines was maintained in a 5% CO_2_ incubator at 37 °C and were routinely tested to exclude *Mycoplasma* contamination.

The CRISPR 3×sgRNA/Cas9 all-in-one expression clone targeting TIGAR (NM_020375.2) (catalogue number: HCP215394-CG04-3) and scrambled sgRNA control for pCRISPR-CG04 (catalogue number: CCPCTR01-CG04-B) were purchased from GeneCopoeia (Rockville, MD). The plasmid DNAs were transformed and amplified in Mix & Go Competent Cells-Strain HB 101 (Zymo Research, catalog no. T3013, Irvine, CA) and purified using PowerPrep HP Plasmid Maxiprep kits with prefilters (Origene, Rockville, MD). The SH-SY5Y neuroblastoma cells were transfected with 10 ug of plasmid DNAs when 60% confluence in 100-mm dishes using transfection reagents GenJet II (SignaGen Laboratories, Rockville, MD per the manufacturer’s instructions). The transfected cells were in culture for 48 hours and were collected in 500 μl of MACS buffer (PBS containing EDTA) containing 1% fetal bovine serum in Falcon 5 ml polystyrene round-bottom tube filtered with cell-strainer cap. The GFP positive cells were selected using FACS GFP sorting and were cultured and proliferated in 10% FBS DMEM. The cell descent was collected and cell lysate was immunoblotted with a TIGAR antibody.

### Method details

#### In vivo ^18^O isotopic labeling of cellular phosphoryls of quardricep white muscle of the mice

Wildtype (WT) and TIGAR knockout (TKO) male mice (n= 3-4 per condition) were at room temperature or shifted to 4°C for 1 h and then given an oral gavage of 0.3 ml pure H_2_^18^O for 10 min at 4°C and an IP injection of 1 ml pure H_2_^18^O for another 10 min at 4° C. The quadricep muscles were freeze-clamped with liquid N_2_ and extracts prepared for mass spectroscopic analyses for identification of the position of ^18^O into the phosphates of ATP, that can be extrapolated to ADP and AMP (Nemutlu, Gupta et al. 2015)

#### Purification and Isotopic Analysis of ^18^O-Labeled Cellular Phosphoryls

Quadricep muscle were freeze-clamped and pulverized in mortar with liquid nitrogen, and extracted in 80% methanol solution. The samples were centrifuged at 10,500 rpm for 10 minutes at 4°C to precipitate proteins. The supernatants were transferred to a LCMS sampling vial pending for injection. Metabolites separation was performed on a ZIC-pHILIC column (Merk). Data was collected with ABSciex 6500+ QTrap MS/MS with a Multiple Reaction Monitoring (MRM) mode. The ions monitored were listed in as below:

| Metabolites | Isotope Labels | Q1 | Q3 |
| --- | --- | --- | --- |
| AMP | M0 | 346 | 79 |
| AMP | M2 | 348 | 81 |
| AMP | M4 | 250 | 83 |
| ADP | M0 | 426 | 79 |
| ADP | M2 | 428 | 81 |
| ADP | M4 | 430 | 83 |
| ADP | M0 | 426 | 159 |
| ADP | M2 | 428 | 161 |
| ADP | M4 | 430 | 163 |
| ADP | M6 | 432 | 165 |
| ATP | M0 | 506 | 79 |
| ATP | M2 | 508 | 81 |
| ATP | M4 | 510 | 83 |
| ATP | M0 | 506 | 159 |
| ATP | M2 | 508 | 161 |
| ATP | M4 | 510 | 163 |
| ATP | M6 | 512 | 165 |
| ATP | M0 | 506 | 239 |
| ATP | M2 | 508 | 241 |
| ATP | M4 | 510 | 243 |
| ATP | M6 | 512 | 245 |
| CP | M0 | 210 | 79 |
| CP | M2 | 212 | 81 |
| CP | M4 | 214 | 83 |

The ^18^O concentration in mouse body was determined with GC-MS according to the MMPC protocol: (https://www.mmpc.org/shared/document.aspx?id=275&docType=Protocol).

#### Stable isotope enrichment and acetylcholine analyses

The SH-SY5Y neuroblastoma cells were cultured in 10 cm dishes until confluent and the culture media was removed, and the cells were washed with DMEM medium (Biological Industries, Boston, catalogue number 01-057-1A, without glucose, glutamine, and pyruvate). Ten ml of DMEM containing 10 mM D-glucose-1,2-^13^C_2_ (Sigma-Aldrich 453188-1G) and 10 mM D-glucose, 1 mM glutamine, without pyruvate was added to the cultured cells and medium was collected at 4, 8, and 24-hour time points. The medium was centrifuged at 1,000 g for 20 min at 4°C and stored at −80°C.

Following thawing on ice, the medium was subjected to an ethyl chloroformate (ECF) derivatization according to our developed procedure with minor modification (Xie, Qiu et al. 2007). Briefly, a volume of 700 ul of cell culture media was added to 500 ul of ethanol: pyridine = 4:1, and 100ul of ECF. After a brief vortex, the samples were sonicated for 1 min. The derivatized metabolites were extracted with 300 ul of chloroform and injected 1ul into the GC-MS system in splitless mode (GC-MS, Agilent, USA). Helium was used as carrier gas at a consistent flow of 1 ml/min. The injection temperature was set at 260 ^0^C and metabolites separation was performed on an Agilent DB-17MS column. The oven program started at 60^0^C for 0.5 min, and rose to 100^0^C at a rate of 20^0^C/min, and then to 300^0^C at a rate of 10 ^0^C/min, and then maintained at 300°C for 4.5 min. The data was analyzed with MassHunter Quantitative Analysis software (Agilent, USA).

The m0 glutamate fragment m/z of 128 contains 3 carbons (carbon 2-4) out the 5 carbons, while fragment m/z of 202 contains 4 carbons (carbon 2-5) from the whole glutamate molecule. M1 of glutamate at m/z 129 reflects the presence of ^13^C at the 4^th^ carbon position. This reflects acetyl units originating from glucose->pyruvate entering through pyruvate dehydrogenase for eventual oxidation (glycolytic flux for energy generation). M2 of glutamate m/z 130 reflects the presence of ^13^C at the 2^nd^ and 3^rd^ carbon position, reflects pyruvate entering the oxaloacetate pool or through pyruvate carboxylase (Madhu, Boneski et al. 2020, Silagi, Novais et al. 2020). M2 of glutamate at 204 is the overall TCA cycle flux. Those two ^13^C come from either pyruvate entering from pyruvate carboxylase or pyruvate dehydrogenase. For lactate, the m/z of M0 to M2 for carbon 2 and 3 position will be 145 to 147 (Madhu, Boneski et al. 2020). A pooled quality control (QC) sample was injected six times for coefficient of variation (CV) determination. Metabolites with CVs lower than 30% were included for quantification while loss with CVs greater than 30% were considered as indeterminant. These data were obtained using a Sciex 6500+ QTRAP with ACE PFP and MERCK Zic-pHILIC columns as previously described (Hoshino, Matoba et al. 2012).

#### Acetylcholine enrichment

The cell culture medium samples were thawed on ice and 200 ul was added to 800 ul of acetonitrile plus 10ul of 10ng/ml acetylecholine_d9 (deuterium 9 as an internal control).

The samples were vortex, centrifuged at 14000rpm for 10 min and supernatants were transferred into glass vials. The samples were analyzed with ABSciex 6500+ with ACE PFP column. The MRMs for m0, m1, m2 for acetylcholine and the internal standard are as following table. Data was processed with multiquanta software (ABSciex). The enrichment was calculated after subtracted from the background of the non-labeled treatment samples.

MRM table

|  |  | Q1 mass (Da) | Q3 mass (Da) |
| --- | --- | --- | --- |
| 1 | Acetylcholine | 146.3 | 87.0 |
| 2 | Acetylcholine_m1 | 147.3 | 88.0 |
| 3 | Acetylcholine_m2 | 148.3 | 89.0 |
| 4 | (d9) Acetylcholine | 155.3 | 87.0 |

#### Total RNA extraction and quantitative RT-PCR

Cellular and tissue total RNA was extracted using TRI reagent and Direct-zol RNA MiniPrep Plus kit (Zymo Research, Irvine, CA). First-strand cDNA was synthesized using the SuperScript^TM^ IV VILO^TM^ cDNA synthesis kit (Thermo Fisher Scientific). TaqMan RT-PCR was performed for measurement of mRNA using the ΔΔ*C_t_* method. Gene expression was adjusted by comparison with *Rpl7* expression. The quantitative RT-PCR results were analyzed by RG Manager version 1.2.1 (Applied Biosystems). Primer-probe mixtures for genes of Ucp1, Atp2a1 (SERCA1), Atp2a2 (SERCA2), Sln (sarcolipin), Mrln (myoregulin), Chrna1, Chrnb1, Chrnd, Chrne, and Chrng were listed as below:

| **Gene name** | **Gene name** | **Taqman Assay ID** |
| --- | --- | --- |
| Uncoupling protein 1 (mitochondrial, proton carrier) | Ucp1 | Mm01244861_m1 |
| ATPase, Ca++ transporting, cardiac muscle, fast twitch 1 (SERCA1) | Atp2a1 | Mm01275320_m1 |
| ATPase, Ca++ transporting, cardiac muscle, slow twitch 2 (SERCA2) | Atp2a2 | Mm01201431_m1 |
| Sarcolipin | Sln | Mm00481536_m1 |
| Myoregulin | Mrln | Mm01175781_m1 |
| Cholinergic receptor, nicotinic, alpha polypeptide 1 (muscle) | Chrna1 | Mm00431629_m1 |
| Cholinergic receptor, nicotinic, beta polypeptide 1 (muscle) | Chrnb1 | Mm00680412_m1 |
| Cholinergic receptor, nicotinic, delta polypeptide | Chrnd | Mm00445545_m1 |
| Cholinergic receptor, nicotinic, epsilon polypeptide | Chrne | Mm00437411_m1 |
| Cholinergic receptor, nicotinic, gamma polypeptide | Chrng | Mm00437419_m1 |

Primer-probe mixture for *Rpl7* was customized, and other primer-probe mixtures were obtained from Thermo Fisher Applied Biosystems.

#### Immunoblotting

Commercial primary antibodies were purchased for the detection of TIGAR (AB10545, Millipore/Sigma-Aldrich), ChAT (703789, ThermoFisher Scientific), VAChT (Cat# 24286, ImmunoStar), c-fos (2250S, Cell Signaling; MA5-15055, Invitrogen), phosphor-c-Fos^Ser32^ (5348S, Cell Signaling), UCP1 (ab209483, abcam), vinculin (sc-73614), and β-actin (sc-47778, Santa Cruz). Samples were prepared from culture cells (washed by cold PBS) or tissues by homogenization with a radioimmune precipitation assay lysis buffer (sc-24948, Santa Cruz Biotechnology) containing Halt^TM^ protease and phosphatase inhibitor mixture (catalog no. 78442, Thermo Fisher Scientific), 100 μM MG132, and 100 μM ALLN (EMD Millipore, Darmstadt, Germany) using Ceria stabilized zirconium oxide beads (MidSci, Valley Park, MO). Homogenates were centrifuged for 15 min at 21,000 × *g* at 4 °C, and supernatants were collected for the protein assay using BCA method. Protein samples were separated by SDS-PAGE and transferred to nitrocellulose membrane using iBlot Blotting System (Thermo Fisher Scientific). The immunoblot membrane was blocked with Pierce Protein-Free T20 (TBS) blocking buffer (product no. 37571, Thermo Fisher Scientific) and incubated with the first antibody indicated in the blocking buffer. Blots were washed in TBS with Tween 20 (TBST) and incubated with either IRDye 800CW secondary antibody (LI-COR, Lincoln, NE) or horseradish peroxidase–conjugated secondary antibody in blocking buffer. The Membrane was washed with TBST and visualized either by the Odyssey Imaging System (LI-COR) or enhanced chemiluminescence (ECL) (Thermo Fisher Scientific Pierce) method. ImageJ was used to quantify protein bands on the membrane.

#### Fluorescence imaging of nicotinic acetylcholine receptors clustering at neuromuscular junctions

Whole Extensor Digitorium Longus (EDL) muscle were fixed 30 min with 4% w/v paraformaldehyde, after dissection. After PBS washing 5 min X 3 times, EDL muscle were incubated with 1:1000 dilution of Alexa Fluor™ 488 conjugate α-Bungarotoxin (Invitrogen B13422) for 2 hours at room temperature, and mount onto 35 mm glass bottom dish (MatTek Corporation, Cat# P35G020C.S) with Vectashield Antifade Mounting Medium with DAPI (Cat# H1200, Vector Laboratories, Inc., Burlingame, CA). The neuromuscular junction was visualized using a z-series projection on a confocal microscope (Leica SP8) with 40X objective. Velocity software was used for image analysis.

#### Immunofluorescence imaging of superior cervical ganglion (SCG)

The ChAT^cre^ and chTKO male mice (forteen-week old) were euthanized and the fresh SCG tissues were removed and embedded in optimal cutting temperature compound. The frozen tissue cross-sections (10 μm) were blocked with 3% bovine serum albumin in phosphate-buffered saline (PBS) for 60 min at room temperature. For vesicular acetylcholine transporter (VChAT) and TIGAR immunofluorescence staining, The SCG section was incubated with the mouse monoclonal anti-TIGAR (E-2) Alexa Fluor 488 (sc-166290 AF488, Santa Cruz Biotechnology) and Goat polyclonal anti-VChAT (Cat# 24286, ImmunoStar, Hudson, WI) for 2 hours at room temperature. After washing PBS 5 min X 3 times, Alexa Fluor 594 donkey anti-goat IgG (H+L) (1:1000, Cat# A-11058, ThermoFisher Scientific) was added to the sections for 30 min at room temperature. The slides were mounted cover slips with Vectashield Antifade Mounting Medium with DAPI (Cat# H1200, Vector Laboratories, Inc., Burlingame, CA), and visualized using a z-series projection on a confocal microscope (Leica SP8) with 40X objective.

#### Synchronized measurement of core body temperature, blood pressure, heart beating rate, and electromyography

ChAT^cre^ and chTKO mice were anesthetized with isoflurane and implanted with DSI HD-X11 telemetric transponder probes according to manufacturer’s guidelines, with the blood pressure probe placed in the descending aorta, the EMG leads run subcutaneously to the neck muscles bilaterally, and the core temperature probe placed intraperitoneally. Animals were returned to a standard temperature housing room maintained at 21-22°C on a 12h/12h light dark cycle, lights on at 0700. Mice were individually housed on cob bedding in standard polycarbonate home cages that were placed on telemetric DSI platforms to continuously monitor arterial blood pressure, heart rate, core body temperature, and neck EMG muscle activity from the implanted DSI XD-11 probes using LabChart 8.1 software for Macintosh. After 1 week recovery from surgery, animals’ home cage bedding was removed and replaced by stainless steel mesh flooring 1 inch above the base of the cage. Food but not water was then removed from the home cage, and animals in their home cages on DSI platforms were then placed in a 10’x10’ environmental cold room maintained at 4C, 35% humidity for 1 hour from 10:00-11:00 am, and continuous measurements of heart rate, blood pressure, core temperature and EMG activity continued. At the end of the 1 hour cold challenge, mice in their individual cages were then returned to the standard temperature housing room, and food and bedding were replaced.

#### Mitochondrial oxidative phosphorylation (OXPHOS) efficiency and capacity

Respiration of permeabilized muscle fibers was performed as previously described (Heden, Ryan et al. 2017, Johnson, Ferrara et al. 2018, Heden, Johnson et al. 2019). Briefly, a small portion of freshly dissected red gastrocnemius muscle tissue was placed in 7.23 mM K_2_EGTA, 2.77 mM Ca K_2_EGTA, 20 mM imidazole, 20 mM taurine, 5.7 mM ATP, 14.3 mM phosphocreatine, 6.56 mM MgCl_2_.6H_2_O, and 50 mM K-MES (pH 7.1). Fiber bundles were separated and permeabilized for 30 min at 4°C with saponin (30 μg/ml) and immediately washed in 105 mM K-MES, 30 mM KCl, 10 mM K_2_HPO_4_, 5 mM MgCl_2_.6H_2_O, BSA (0.5 mg/ml), and 1 mM EGTA (pH 7.4) for 15 min. After washing, high-resolution respiration rates were measured using an OROBOROS Oxygraph-2k.

#### Muscle contraction/relaxation

The extensor digitorum longus muscle was dissected and tied into the Horizontal Tissue Bath System from Aurora Scientific Inc. as previously described (Ferrara, Verkerke et al. 2018). The muscles were stimulated with a 20-V twitch train and stretched until optimal length for force production was reached. Muscles were then allowed to equilibrate for 5 min. For force-frequency curves, muscles were stimulated with frequencies ranging from 10 to 200 Hz (0.1-ms pulse, 330-ms train, and 2 min between trains). Forces produced by electrically stimulated muscle contractions were recorded in real time via a force transducer (model 400A, Aurora Scientific Inc.). Specific force was calculated using cross-sectional area of the muscle tissue (millinewton per square millimeter), as estimated from the weight and length of the muscle.

#### Gastrocnemius muscle acetylcholine assay

Fresh gastrocnemius muscle was collected and snap-frozen in liquid nitrogen and the whole piece of the muscle was ground to powder using a mortar-pestle in liquid nitrogen. The muscle powder was then homogenized in PBS (pH 7.4) using Ceria stabilized zirconium oxide beads (MidSci, Valley Park, MO) and the homogenate was centrifuged for 20 minutes at 3,000 rpm at 4°C degree. The supernatant was collected and the protein assay was performed using the Pierce BCA Protein Assay kit (Cat#23225, ThermoFisher Scientific). The supernatants from different samples were adjusted to the same protein concentration of protein and the same volume of the supernatants were subjected to acetylcholine assay using QuickDetectTM Acetylcholine (ACh) (Mouse) ELISA Kit (Cat#E4453-100, Biovision, Milpitas, CA).

### Quantification and statistical analyses

The number of independent experimental replications and the average with standard deviation are provided in the figure legends. The data were analyzed by one-way analysis of variance using Prism version 8 software (GraphPad Software, Inc., La Jolla, CA) for comparison of multiple groups or unpaired *t* test for two groups. The statistical analyses were made at significance levels as follows: ns, not statistically significant; \**p* < 0.05; \*\**p* < 0.01; \*\*\**p* < 0.001; \*\*\*\**p* < 0.0001.

## Supplemental figures legends

**Figure S1:**
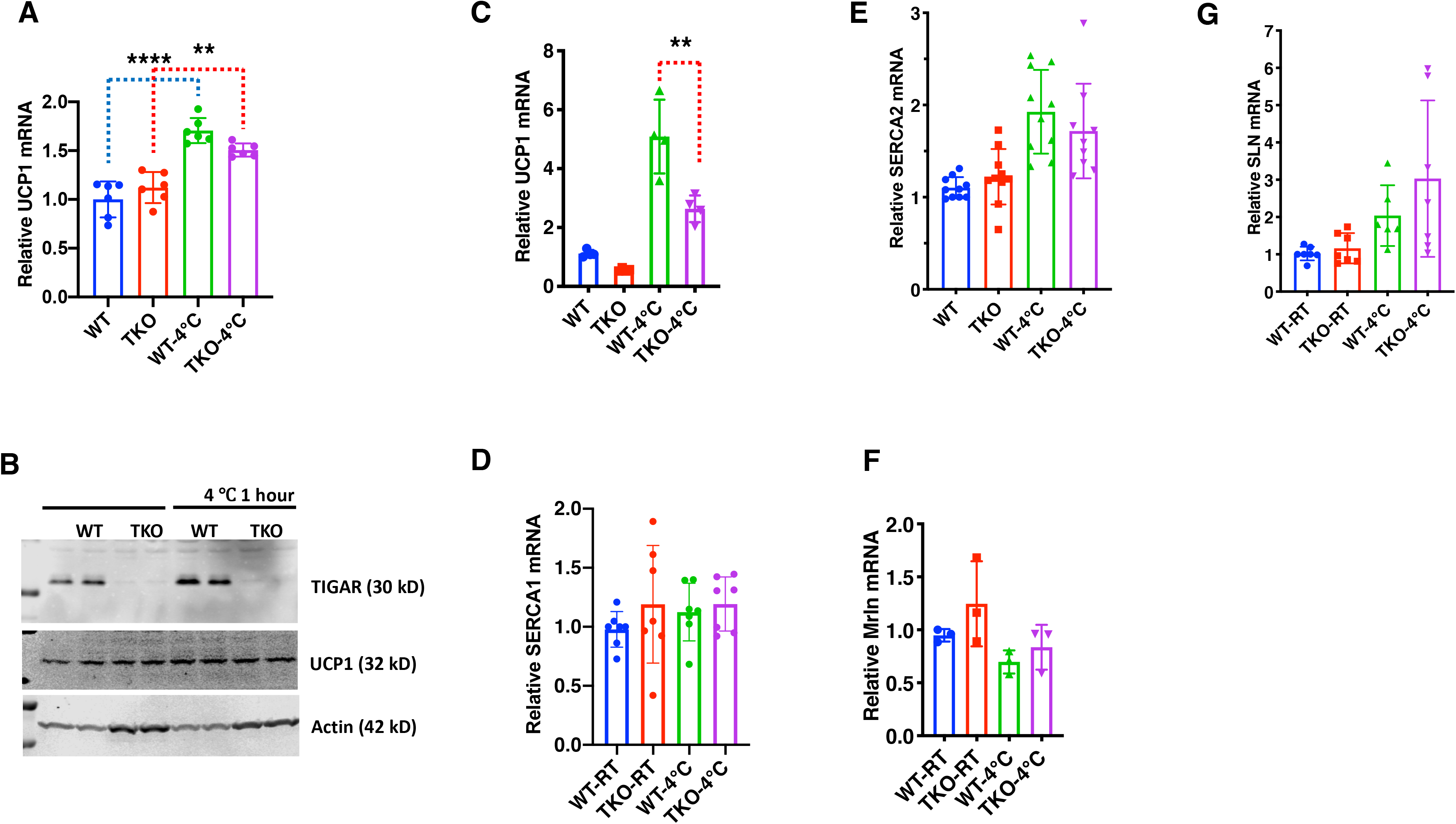
The TKO protection against hypothermia is independent of UCP1 and SERCA expression. A, Relative UCP1 mRNA levels in interscapular brown adipose tissue from wildtype (WT) ad TKO male (n=6) mice maintained at ambient temperature and after 1 hour 4°C exposure. B, Representative TIGAR, UCP1 and actin protein immunoblot of interscapular brown adipose tissue from two independent wildtype (WT) and TKO male mice maintained at ambient temperature or after 1 hour 4°C exposure. C, Relative UCP1 mRNA levels in inguinal adipose tissue from WT and TKO male mice (n= 4-10) at ambient temperature or after 2-hour 4°C exposure. Please note that Ct values for brown adipose tissue was approximately 18, whereas the Ct values for the inguinal white adipose tissue was in the range of 33. Relative mRNA levels of D, SERAC1, E, SERCA2, F, myoregulin (Mrln) and G, sarcolipin (SLN) in the gastrocnemius muscle from WT and TKO mice at ambient temperature or after a 1 hour 4°C exposure. Statistical analyses were performed using Prism8 and data presented is the mean ± S.D. with two-way ANOVA multiple comparison. *p* < 0.01, **; p<0.0001, ****.

**Figure S2:**
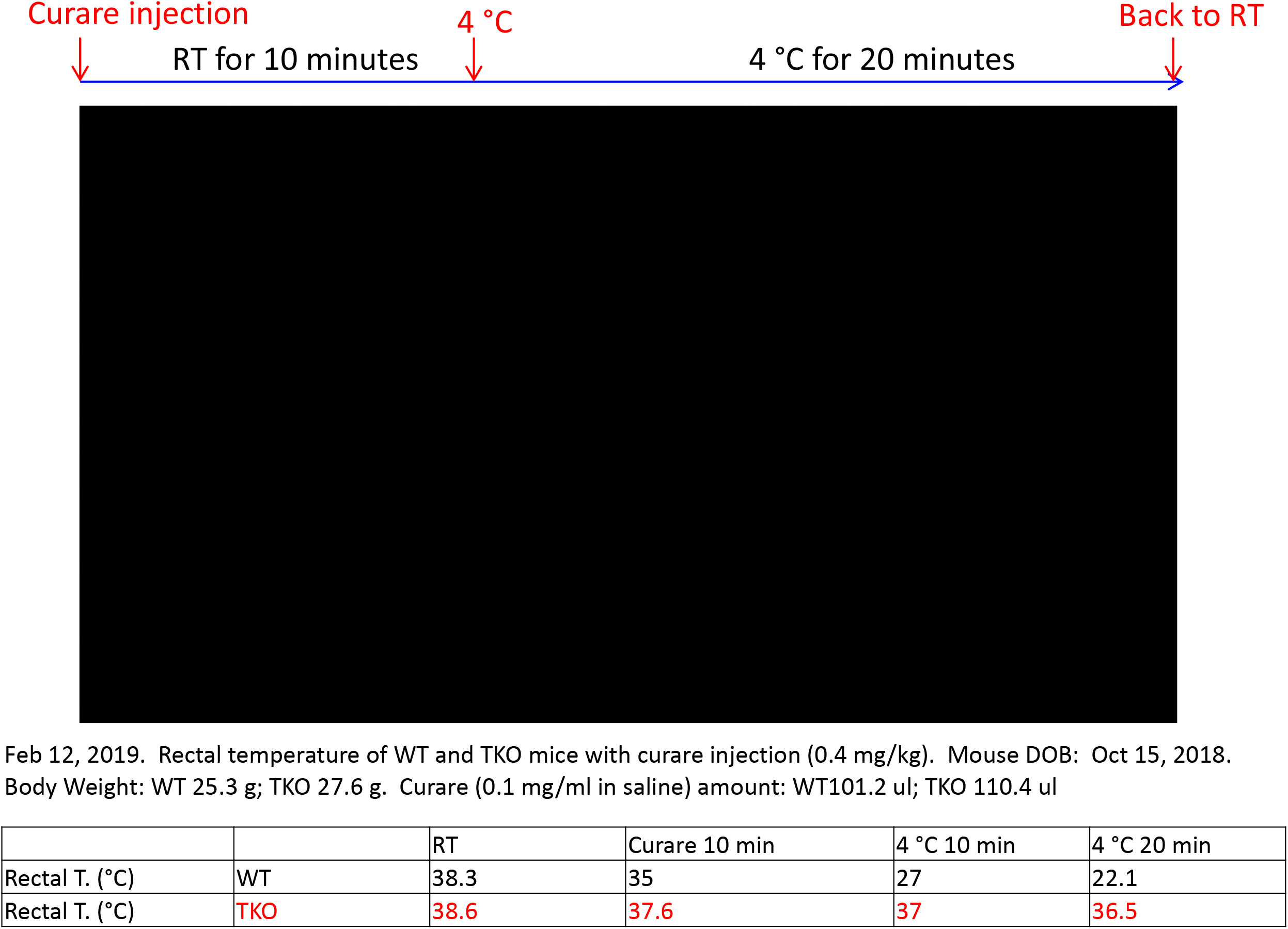
TKO mice are resistant to the paralytic effects of tubocurare. Representative video clip of wildtype (WT) and whole-body TIGAR knockout (TKO) male mice behavior under ambient temperature and at 4°C following and intraperitoneal injection of tubocurare (0.4mg/kg body weight).

**Figure S3:**
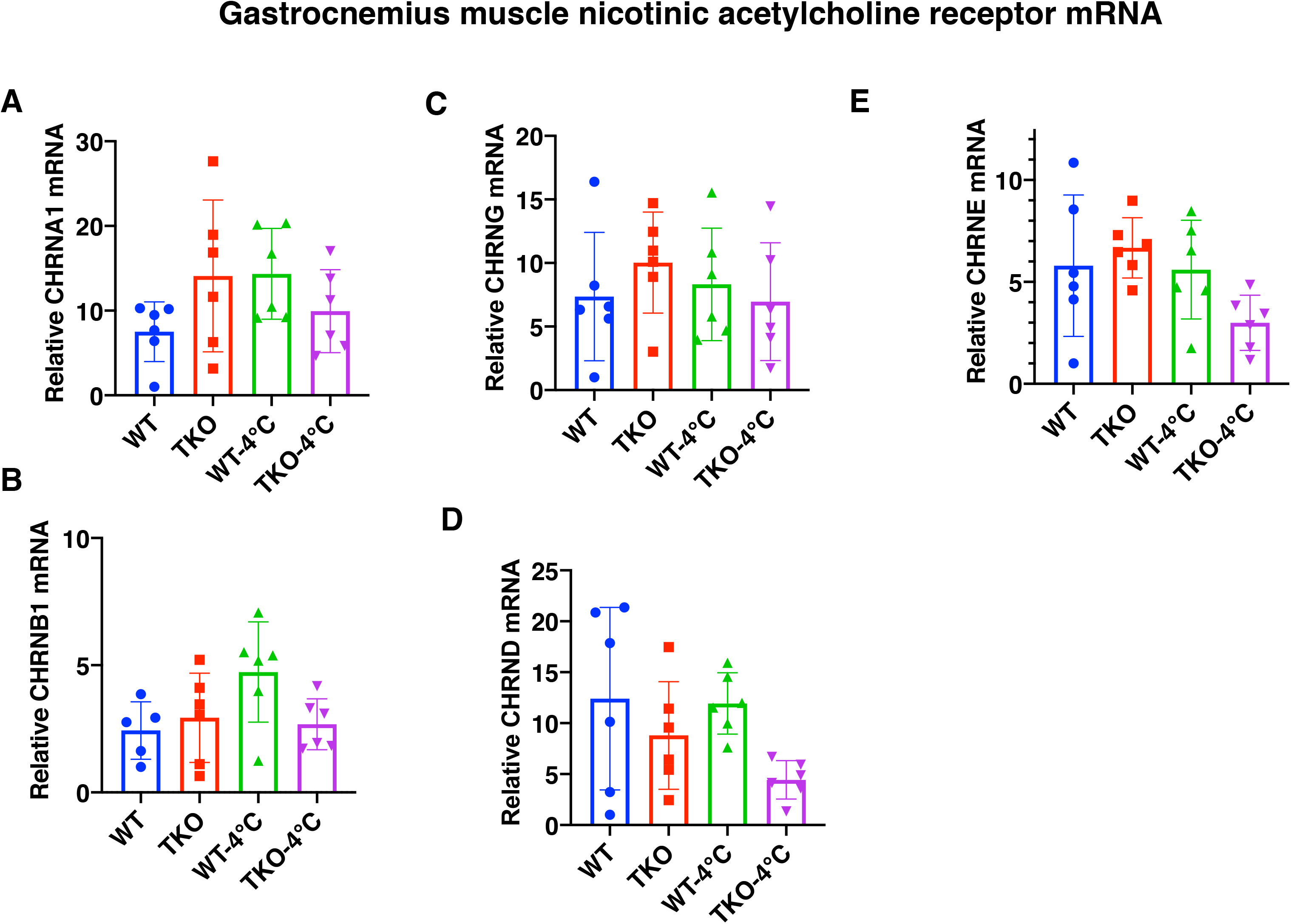
Changes in the expression of skeletal muscle nicotinic acetylcholine receptor subunits does not account for the differential sensitivity to tubocurare. Relative mRNA levels of the nicotinic acetylcholine receptor subunits A, α1, B, β1, C, γ D, δ, and E, ε in the gastrocnemius muscle from male (n=5-6) wildtype (WT) and whole-body TIGAR knockout (TKO) mice at ambient temperature or after one-hour 4°C exposure.

**Figure S4:**
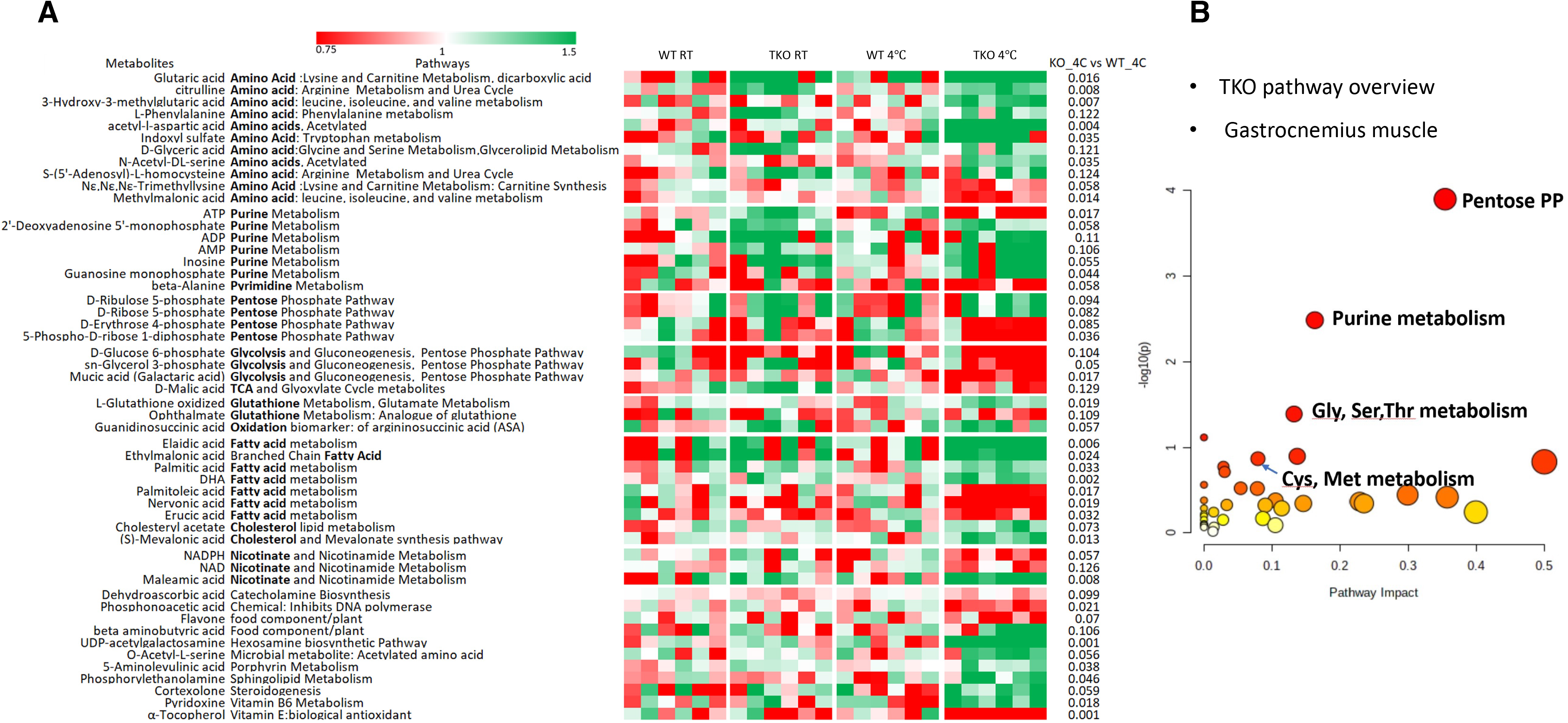
Skeletal muscle of TKO mice at 4°C display increased pentose phosphate pathway, purine nucleotide cycle, and amino acid utilization pathways. Wildtype (WT) and whole-body TIGAR knockout (TKO) male mice (n=6) were maintained at room temperature (RT) or shifted to 4°C for 1 hour. a, Quadriceps white skeletal muscle was isolated and extracts were subjected to widely targeted (MRM) small metabolite profiling using an ABSciex 6500+ QTRAP with ACE PFP and MERCK Zic-pHILIC columns as described in Method and Materials. The heatmap shows the metabolites/pathways differentially identified with corresponding p values. B, Matched pathways, determined from MetaboAnalyst 5.0, which integrates enrichment analysis (metabolite over-representation more than expected by chance) and pathway impact (reflecting centrality of the metabolite in a topological network).

**Figure S5:**
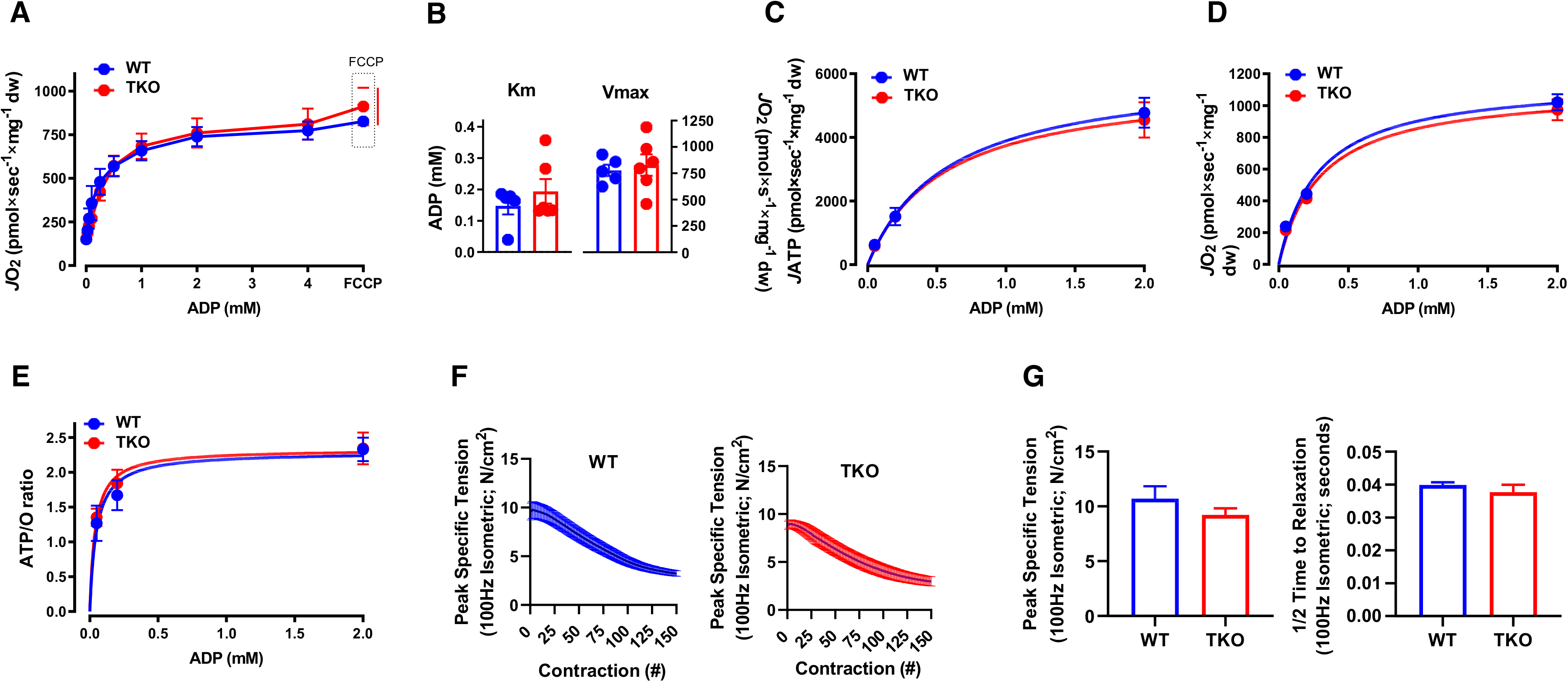
TIGAR deficiency has no significant effect on intrinsic skeletal muscle mitochondrial oxidative phosphorylation activity or ATP production. Permeabilized skeletal muscle fiber bundles were prepared from red portions of the gastrocnemius muscle from wildtype (WT) and TKO mice immediately after 1 h cold exposure. A, Oxygen consumption rate (*J*O_2_) as a function of ADP concentration. B, The Km and Vmax for ADP-stimulatory respiratory kinetics. C, Rates of ATP production (*J*ATP) and D, oxygen consumption (*J*O_2_) measured simultaneously under clamped submaximal and maximal ADP-demand states. E, ATP/O efficiency ratios. F, Force-frequency of the extensor digitorum longus muscle isolated from WT and TKO mice. G, Peak specific tension and time to one-half relaxation of the extensor digitorum longus muscle from WT and TKO mice.

**Figure S6:**
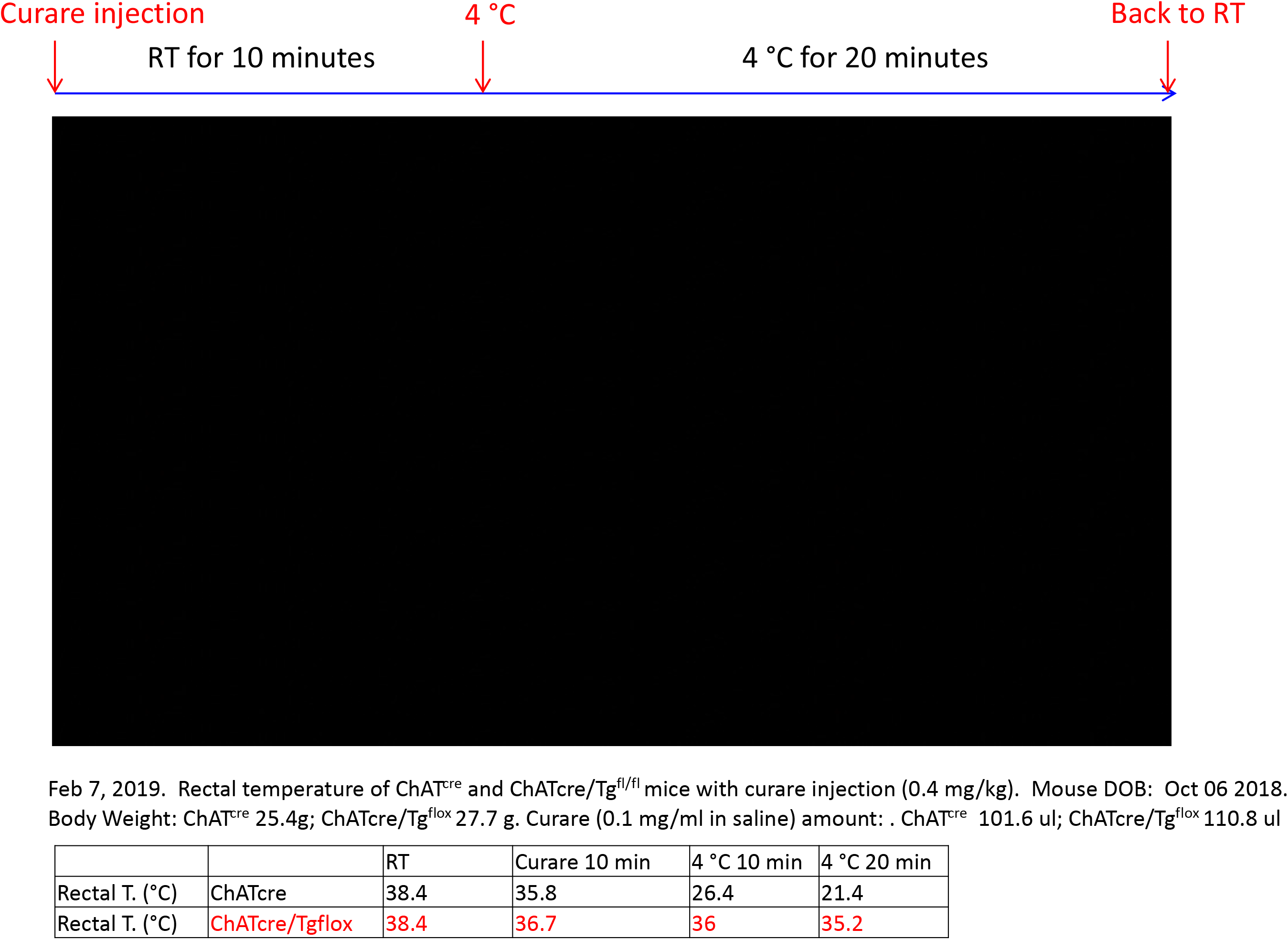
Cholinergic neuron specific TIGAR knockout (chTKO) mice are resistant to the paralytic effects of tubocurare. Representative video clip of control (ChAT^cre^) and chTKO male mice behavior under ambient temperature and at 4°C following and intraperitoneal injection of tubocurare (0.4mg/kg body weight).

**Figure S7:**
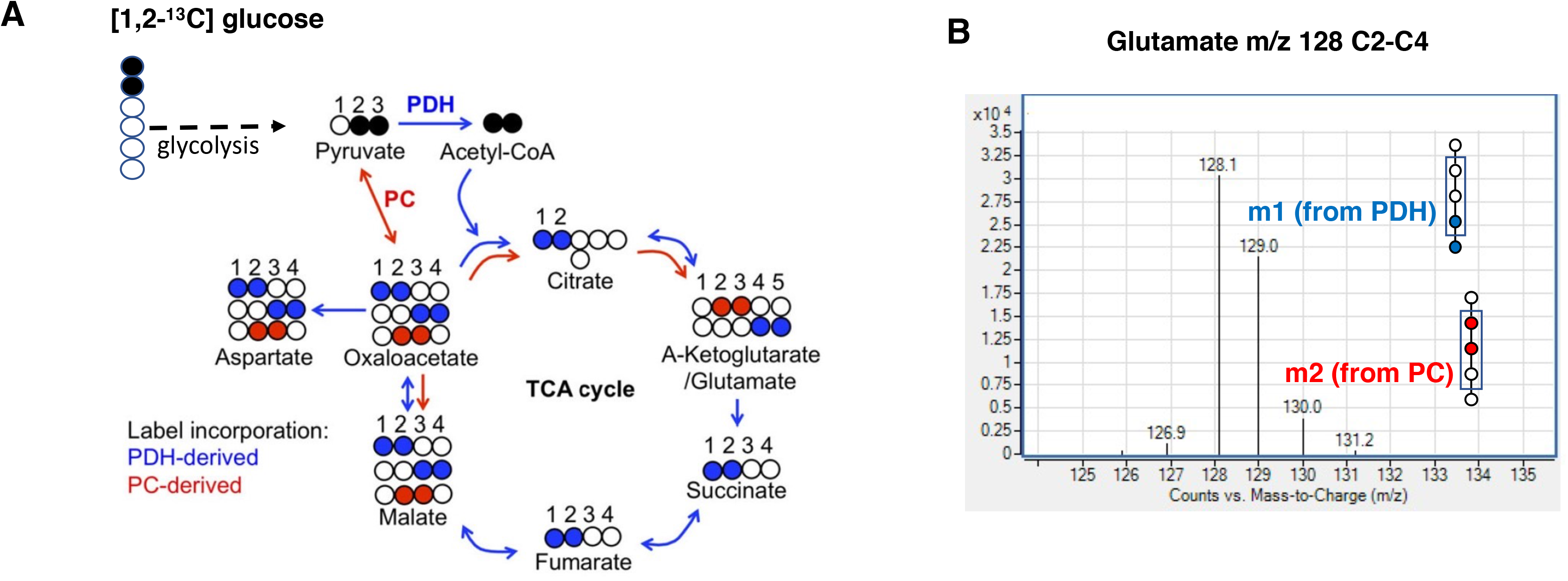
Summation of flux results through glycolysis and the TCA cycle using [1,2]-^13^C-glucose by assessing glutamate isotopomers. A, [1,2]-^13^C-glucose becomes [2,3]-^13^C pyruvate if it is directly metabolized through the glycolytic pathway. [2,3]-^13^C-pyruvate can enter the TCA cycle through pyruvate dehydrogenase (PDH) or pyruvate carboxylase (PC). Blue dots indicate ^13^C carbons from glucose entered TCA cycle via PDH. Red dots indicate ^13^C carbons from glucose entered TCA cycle via pyruvate carboxylase (PC). B, Representative spectra of the glutamate C2-C4 fragments from TIGAR knockout SH-SY5Y cells labeled with D-glucose-[1,2]-^13^C_2_. M0 of the C2-C4 glutamate fragment is m/z 128, m1 of glutamate is m/z 129 (blue) that indicates the presence of ^13^C at the fourth carbon position and m2 glutamate indicates the presence of 13C at the second and third carbon position (red).

